# rG4-seeker enables high-confidence identification of novel and non-canonical rG4 motifs from rG4-seq experiments

**DOI:** 10.1101/2020.02.01.929851

**Authors:** Eugene Yui-Ching Chow, Kaixin Lyu, Chun Kit Kwok, Ting-Fung Chan

## Abstract

We recently developed the rG4-seq method to detect and map *in vitro* RNA G-quadruplex (rG4s) structures on a transcriptome-wide scale. rG4-seq of purified human HeLa RNA has revealed many non-canonical rG4s and the effects adjacent sequences have on rG4 formation. In this study, we aimed to improve the outcomes and false-positive discrimination in rG4-seq experiments using a bioinformatic approach. By establishing connections between rG4-seq library preparation chemistry and the underlying properties of sequencing data, we identified how to mitigate indigenous sampling errors and background noise in rG4-seq. We applied these findings to develop a novel bioinformatics pipeline named rG4-seeker(https://github.com/TF-Chan-Lab/rG4-seeker), which uses tailored noise models to autonomously assess and optimize rG4 detections in a replicate-independent manner. Compared with previous methods, rG4-seeker exhibited better false-positive discrimination and improved sensitivity for non-canonical rG4s. Using rG4-seeker, we identified novel features in rG4 formation that were missed previously. rG4-seeker provides a reliable and sensitive approach for rG4-seq investigations, laying the foundations for further elucidation of rG4 biology.

## INTRODUCTION

Guanine-rich sequences can interact at Watson-Crick and Hoogsteen edges via hydrogen bonds and fold into non-canonical structural motifs termed G-quadruplexes (G4s)^1^. G4s consist of two or more layers of G-quartet planes, which are connected by loop nucleotides^2,3^. G-quartets in G4s are additionally stabilized by monovalent cations, in which Li^+^ < NH4^+^ < Na^+^ < K^+1^. RNA G4s (rG4s) have previously been visualized in both fixed and live cells using G4-specific antibodies and G4-specific fluorescence probes^4–7^. Numerous studies have suggested that rG4s regulate processes such as transcription, RNA splicing, translation and RNA metabolism^8^.

Computational tools have been developed to predict rG4 forming potential in the transcriptome^2,3^. While these computational tools are applicable to any transcriptome of interest, their algorithms are largely based on putative G4 sequence motif searches and GC content, and they are derived from limited *in vitro* experimental datasets^2,3^. However, these low-throughput experimental assays work well on relatively short oligonucleotides^2,3^. We recently developed a high-throughput sequencing method referred as rG4-seq to detect and map *in vitro* rG4 structures on a transcriptome-wide scale^9^. Applying rG4-seq to purified human HeLa RNA revealed that *in vitro* rG4 formation propensity is significantly influenced by the flanking sequence^9^. In addition, many non-canonical rG4 motifs (such as long loops, bulges, and 2-quartet) have been identified^9^, which are challenging to predict using primary RNA sequencing alone.

Previously, we developed a bioinformatic analysis pipeline to identify rG4-induced reverse transcriptase stalling (RTS) sites from a HeLa rG4-seq dataset based on coverage drop signal (CDS) metrics, which measure the local changes in sequencing read coverage in an averaging window 21 nt long^9^. However, evaluation of the flanking nucleotide sequences has revealed approximately 12% of the readouts to be rG4s false positives^9^. Given that four biological replicates and stringent statistical methods have been used, it is unlikely that those false-positive detections were coincidental. We hypothesized that the origins of the false positives relate to the underlying properties of the bioinformatics pipeline and the dataset. Using an alternative RTS metric and statistical methods, we identified how to mitigate indigenous sampling errors and background noise in rG4-seq, and developed a new bioinformatics pipeline, rG4-seeker that uses tailored noise models to autonomously assess and optimize the detection of rG4-induced RTS sites. Notably, rG4-seeker revealed previously unidentified features in the HeLa rG4-seq dataset. These novel features were also experimentally validated in this study.

## RESULTS

### Ratio of stalled reads (RSR) metric improves resolution of RTS events

To scrutinize the design of our rG4-seq bioinformatics pipeline, we studied other bioinformatic pipelines designed for analyzing datasets from SHAPE-seq and eCLIP experiments^10,11^. The two techniques shared similarity with rG4-seq in exploiting the RTS phenomenon to detect RNA structures and elements. Interestingly, we found that all pipelines preferred to use read starts, which indicate the nucleotide position where RT terminates as the signal for RTS events.

To utilize read start information in rG4-seq analysis, we normalized the read start counts in the rG4-seq dataset by the read coverage to obtain the RSR metric. RSR indicates the observed frequency of an RTS event at a single nucleotide position (assuming a binomial distribution B(*n*, *p*), where *n* is the read coverage and *p* is the underlying probability of RTS events). Genuine rG4s are expected to induce RTS events at the 3’ end, which lead to single-nucleotide RSR peaks that are significantly higher in rG4-stablizing (K^+^ or K^+^-PDS) conditions than in rG4-non-stabilizing (Li^+^) conditions.

To compare the newly proposed RSR metric and the original CDS metric, we re-analyzed the HeLa rG4-seq dataset generated in our previous study^9^ (see methods). We found that the RSR metric improved the resolution for closely adjacent RTS events and weak RTS events (Figure 1A). In agreement with the findings from previous polyacrylamide gel electrophoresis (PAGE)-based RTS assays^9,12^, the rG4-induced RSR peaks was detected specifically at or adjacent to the 3’ most guanine residues (Figure 1A). Moreover, among the false-positive RTS sites that were assigned to the “Others” category in our previous study, a considerable proportion of them did not associate with any statistically significant RSR peaks (binomial test, *p* <0.05) (Figure 1B). These initial findings suggested that given a proper statistical testing scheme, the RSR metric effectively distinguished true positives from false-positive RTS events.

**Figure 1.**
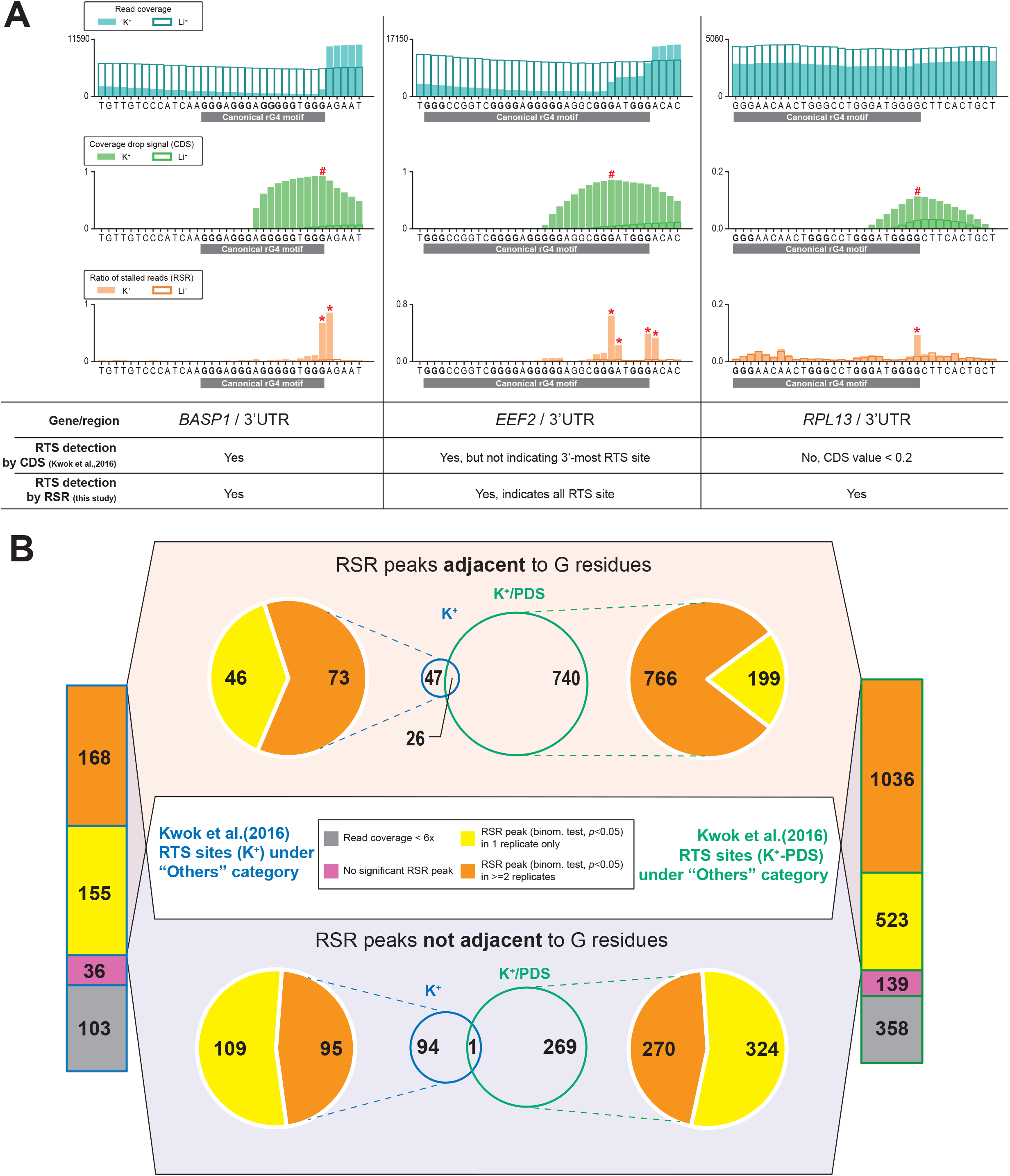
Ratio of stalled reads (RSR) metric improves resolution and specificity for detecting RTS events. (a) Showcases comparing proposed ratio of stalled reads (RSR) metric with coverage drop signal (CDS) metric used to resolve RNA G-quadruplex (rG4)-induced reverse transcriptase stalling (RTS) events.

- In the 1^st^ showcase at the *BASP1* gene, the RTS event associated with the canonical rG4 was detected by both CDS and RSR. While both metrics indicated a signal peak at the 3’ end of the rG4 motif, RSR produced a sharper and narrower peak.
- In the 2^nd^ showcase at the *EEF2* gene, the RTS event was detected by both CDS and RSR. This rG4 motif exhibited 5 G tracts and utilized either the 4^th^ or 5^th^ G tract as the 3’-most G tract. The RSR metric offered higher resolution and indicated both RTS sites corresponding to the 4^th^/5^th^ G tract with two separate peaks
- In the 3^rd^ showcase at the *RPL13* gene, the RTS event was detected by RSR but not CDS owing to low RTS effect strength as nucleotide positions of CDS <0.2 were removed to eliminate low-confidence data points. Compared to CDS, the RSR metric offered better resolution for closely adjacent RTS events or weak RTS events. (b) Re-analysis of previously reported RTS sites under the “Others” category using the RSR metric. RTS sites under the “Others” category are considered false positives and should be minimized in rG4-seq analysis. The proposed scheme rejected most of these RTS sites by criteria of insufficient read coverage (<6x), absence of associated reproducible RSR peaks, or disagreements between rG4-seq (K^+^) and rG4-seq (K^+^-PDS) experiments. Meanwhile, 27 “Others” RTS sites met all detection criteria, where some were genuine rG4s incorrectly classified as false positives.

To better account for the under-sampling errors of RSR measurements among low-abundance transcripts (Figure 2A), we derived a modified statistical testing scheme named “minimum ΔRSR” that considered read coverages in datasets from both rG4-stabilizing and rG4-non-stabilizing conditions (Figure 2B). Based on the re-analysis of the “Others” RTS sites, we found that the minimum ΔRSR scheme out-performed the conventional binomial test and rejected most false positive sites (Figure 1B, Figure 2C), where a majority of them had a read coverage of <30x (Table 1). Meanwhile, the minimum ΔRSR scheme also identified 21 “Others” RTS sites that were not genuine false positives. These sites were detected in both K^+^ and K^+^-PDS conditions and occurred adjacent to 1 or more GGG motifs that resembled G-tracts (Figure 2C, Supplementary Table S1).

**Table 1.** False-positive RTS site detection are associated with low read coverage.

| Properties of “Others” RTS sites (K<sup>+</sup>) |  |  |  |
| --- | --- | --- | --- |
| Has reproducible RSR peaks | Yes | Yes | Yes |
| Adjacent to G residues | No | Yes | Yes |
| Overlaps with “Others” RTS sites (K<sup>+</sup>-PDS) | No | No | Yes |
| Total no. of RTS sites | 94 (100%) | 47 (100%) | 25 (100%) |
| Read coverage < 10x | 29 (31%) | 18 (38%) | 2 (8%) |
| 10x ≤ Read coverage < 30x | 53 (56%) | 24 (51%) | 0 (0%) |
| 30x ≤ Read coverage < 50x | 6 (6%) | 4 (9%) | 10 (38%) |
| Read coverage ≥ 50x | 6 (6%) | 1 (2%) | 14 (54%) |

Properties of “Others” RTS sites (K<sup>+</sup>) that were found to have reproducible RSR peaks in ≥2 replicates (binomial test, $p < 0.05$ ). For RTS sites (K<sup>+</sup>) that were either not adjacent to G residues or not overlapping with “Others” RTS sites (K<sup>+</sup>-PDS), vast majority of them have read coverage < 30x. In contrast, for the 25 candidates that overlapped with RTS sites (K<sup>+</sup>-PDS), more than half of them have read coverage ≥50x.

**Figure 2.**
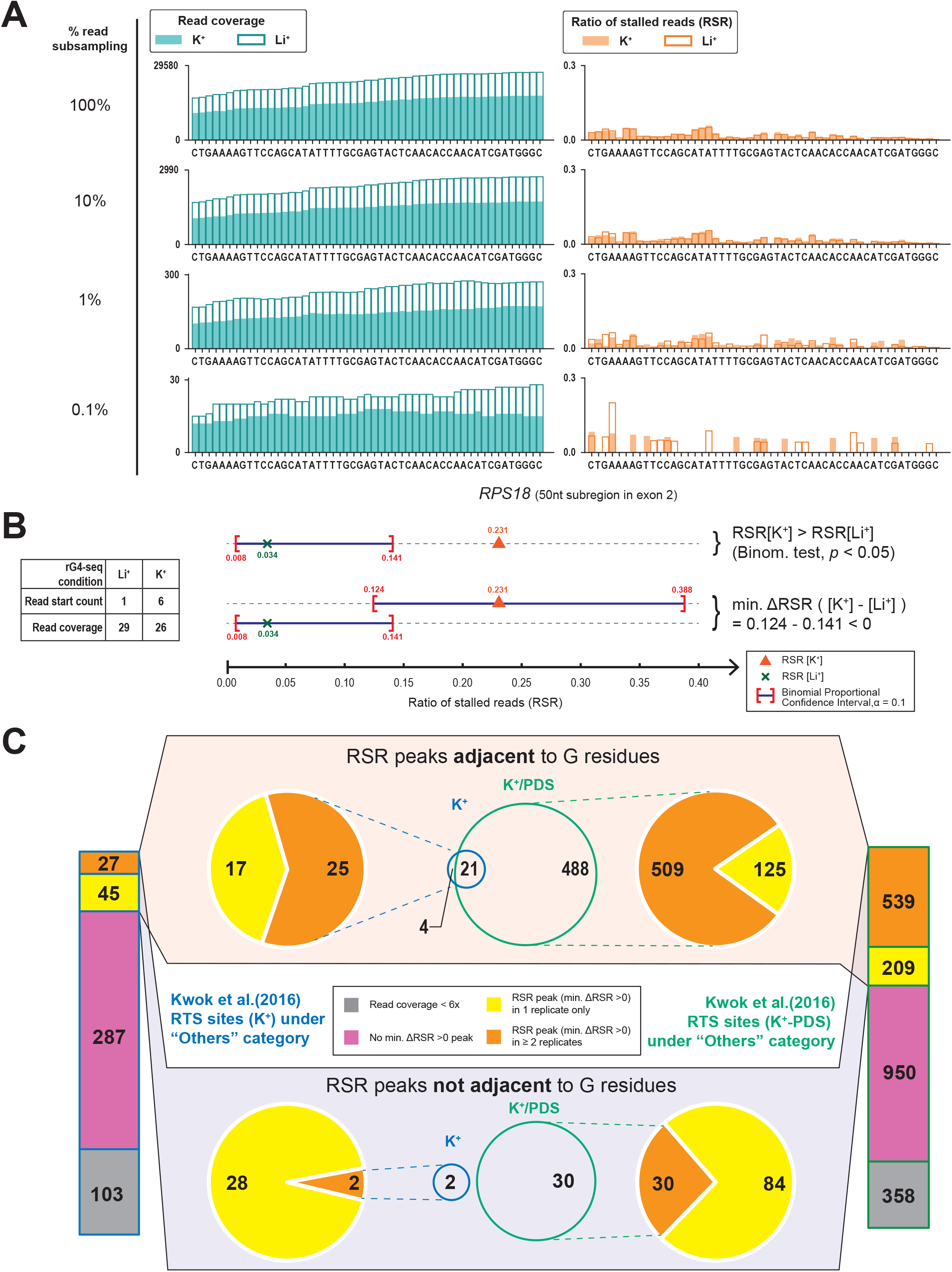
Minimum ΔRSR scheme out-performs binomial test in RTS event detection at lower read coverage. (a) The effects of low read coverage on ratio of stalled reads (RSR) signal simulated by exponential read subsampling, demonstrated by a 50 nt non-rG4-harbouring region on the *RPS18* gene. At the original read coverage, an equivalent RSR signal was recorded by rG4-seq at K^+^ (rG4-stabilizing) and Li^+^ (rG4-non-stabilizing) conditions without statistically significant differences (binomial test), suggesting that the underlying RT stalling probabilities did not differ between the two conditions. However, the shape of the original RSR signal was distorted and the similarity of RSR signal between the two conditions was lost at reduced read coverage (~300x and ~30x) when under-sampling error affected the RSR measurements. (b) Comparison between the binomial test and proposed minimum ΔRSR metric scheme. A marginal case at low read coverage (~30x) extracted from a non-rG4-harboring region in rG4-seq dataset is shown. Minimum ΔRSR metric scheme additionally addresses sampling error of RSR[K^+^] measurement with binomial proportional confidence interval. (c) Re-analysis of previously reported RTS sites under the “Others” category using minimum ΔRSR scheme. RTS sites under the “Others” category are considered false positives and should be minimized in rG4-seq analysis. The minimum ΔRSR scheme out-performed the binomial test in rejecting more “Others” RTS sites and reported less reproducible RSR peaks non-adjacent to G residues. Meanwhile, 21 “Others” RTS sites met all detection criteria and were adjacent to G residues.

Taken together, our findings suggest that the RSR metric and minimum ΔRSR scheme are more reliable in identifying RTS sites from rG4-seq datasets compared with the CDS metric developed previously. Moreover, given the improved resolution of the RSR metric, which guaranteed rG4-induced RTS site detection only at 3’ end of the G-tracts, we could develop a new sequence-based filtering scheme in rG4-seq analysis that would limit RTS detection to the loci where G-tract formation was possible.

### Transcriptome-wide analysis reveals prevalent background noise in rG4-seq data

To apply the RSR metric and minimum ΔRSR scheme in a transcriptome-wide scenario, we next computed statistically significant, single-nucleotide RSR peaks (minimum ΔRSR >0) within the annotated regions of the transcriptome. We found that the RSR peaks of higher minimum ΔRSR values (indicating stronger RTS effect strength) were present only for the RSR[rG4-stabilzing] > RSR[rG4-non-stabilizing] scenario. By overlaying the RSR peaks associated with the canonical/G3L_1-7_ rG4s reported in our previous study, we found that high minimum ΔRSR values were indeed associated with RTS events of genuine rG4s (Figure 3A).

**Figure 3.**
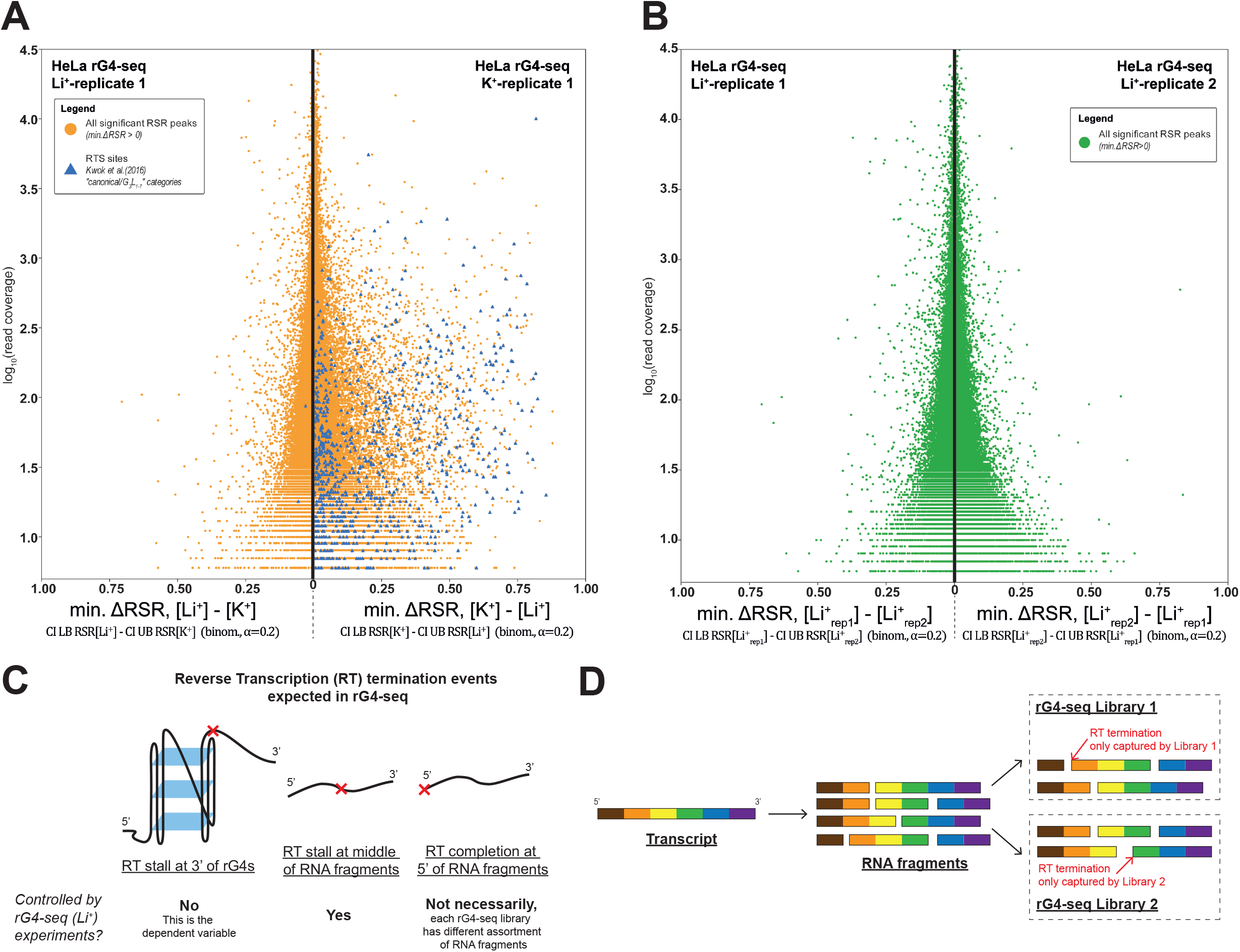
False-positive RTS detections originate from RNA fragmentation-associated background noise. (a) Transcriptome-wide significant RSR peaks (minimum ΔRSR >0) detected from replicate 1 of HeLa rG4-seq dataset (pairwise comparison between K^+^/Li^+^ condition). Minimum ΔRSR values (indicating RTS effect strength) of RSR peaks were plotted against read coverage (logarithmic scale). Each datapoint corresponds to 1 RSR peak at 1 single-nucleotide genomic locus. RSR peaks coinciding with RTS sites in canonical/G3L1-7 reported by Kwok et al. (2016) are highlighted. (b) Transcriptome-wide significant RSR peaks (minimum ΔRSR >0) detected from a pairwise comparison between replicates 1 and 2 of the Li^+^ HeLa rG4-seq dataset. Comparison between replicates of identical conditions implies that all detected RSR peaks originated from experimental variations. (c) Summary of different reverse transcription (RT) termination events expected in rG4-seq that would generate RSR signals, and whether these events were controlled by rG4-seq (Li^+^) experiments. (d) Illustration of noise introduced to rG4-seq sequencing data by the RNA fragmentation process. Each rG4-seq library receives an unidentical assortment of RNA fragments, which causes discrepancies in the RT termination events at 5’ of the RNA fragments between libraries.

We found that the total number of RSR peaks detected greatly exceeded our previous estimation of genuine rG4 numbers in the transcriptome (hundreds of thousands versus thousands), and the majority of peaks had lower minimum ΔRSR values and were distributed in a pyramidal region at the center of the plot (Supplementary Figure S2, S3). In contrast, for the negative controls where RSR[rG4-stabilzing] < RSR[rG4-non-stabilizing], the RSR peaks were exclusively distributed within the pyramidal region. We reasoned that most of these RSR peaks of lower minimum ΔRSR originated from background noise in the rG4-seq.

To further explore this phenomenon, decoy experiments were conducted to compare the six pairwise combinations of the four Li^+^ condition replicates. We hypothesized that any significant RSR peaks (minimum ΔRSR >0) identified would originate from experimental variations between replicate experiments and background noise of rG4-seq. Strikingly, we found that the pyramidal distribution of RSR peaks in the center of the plot was reproduced in these decoy experiments, indicating the presence of background noise and that magnitude was inversely proportional to read coverage. In contrast, RSR peaks of higher minimum ΔRSR values were not detected among the decoys (Figure 3B, Supplementary Figure S4). Moreover, the distribution of decoy RSR peaks was found to be asymmetric, where the total number of detected RSR peaks varied between 75,000 to 200,000 depending on the combination of replicates compared (Figure 3B, Supplementary Figure S4). These observations suggested that there could be benign intrinsic differences between individual rG4-seq datasets independent of the experimental conditions, which generated the weak background RSR peaks by coincidence.

### Modelling of background noise associated with RNA fragmentation process

By revisiting the chemistry of rG4-seq, we inferred that the background RSR peaks were likely to be associated with noise introduced during the random RNA fragmentation procedure:

- Apart from the read stop originating from RT stalling (premature termination), read stops were also recorded when reverse transcription completed at the 5’ end of RNA fragments. These two independent events both contributed to the RSR measurements and are indistinguishable (Figure 3C).
- In each rG4-seq replicate experiment, the input RNA sample was first randomly fragmented and then aliquoted into 3 equal portions for preparing the K^+^, Li^+^ and K^+^-PDS libraries. As each library received a different “cocktail” of RNA fragments, the distribution of 5’ ends of the fragments could not be assumed to be equal between replicate rG4-seq libraries (Figure 3D).
- When one rG4-seq library received more fragments that started at a certain nucleotide position than another library, the event could be translated as an RSR peak (Figure 3D).

The pyramidal distribution of the weak background RSR peaks indicated that their minimum ΔRSR values were bound by a probabilistic upper limit. Importantly, genuine rG4s generated RSR peaks with minimum ΔRSR values above this probabilistic upper limit, because the pyramidal region did not overlap with the stronger RSR peaks associated with canonical/G3L_1-7_ rG4s. These findings prompt us to establish a statistical model to estimate this upper limit by assuming a worst-case scenario of behavior of fragmentation-associated noise. This model excluded weak RSR peaks that were likely to be associated with fragmentation noise.

### New RTS site calling workflow eases replicate requirements and out-performs prior methods

Based on the minimum ΔRSR metric scheme, sequence-based filtering scheme and newly derived fragmentation-associated noise model, *ab initio* RSR peak identification and RTS site calling were conducted independently for each rG4-seq replicate dataset at a target FDR of ≤1.5% (Figure 4A, B, Supplementary Figure S5, S6) (see methods). The RTS sites were then classified into six categories (canonical/G3L_1-7_, long loops, bulges, 2-quartet, G≥40% and Others) based on their adjacent sequences and according to the scheme described in our previous study^9^.

**Figure 4.**
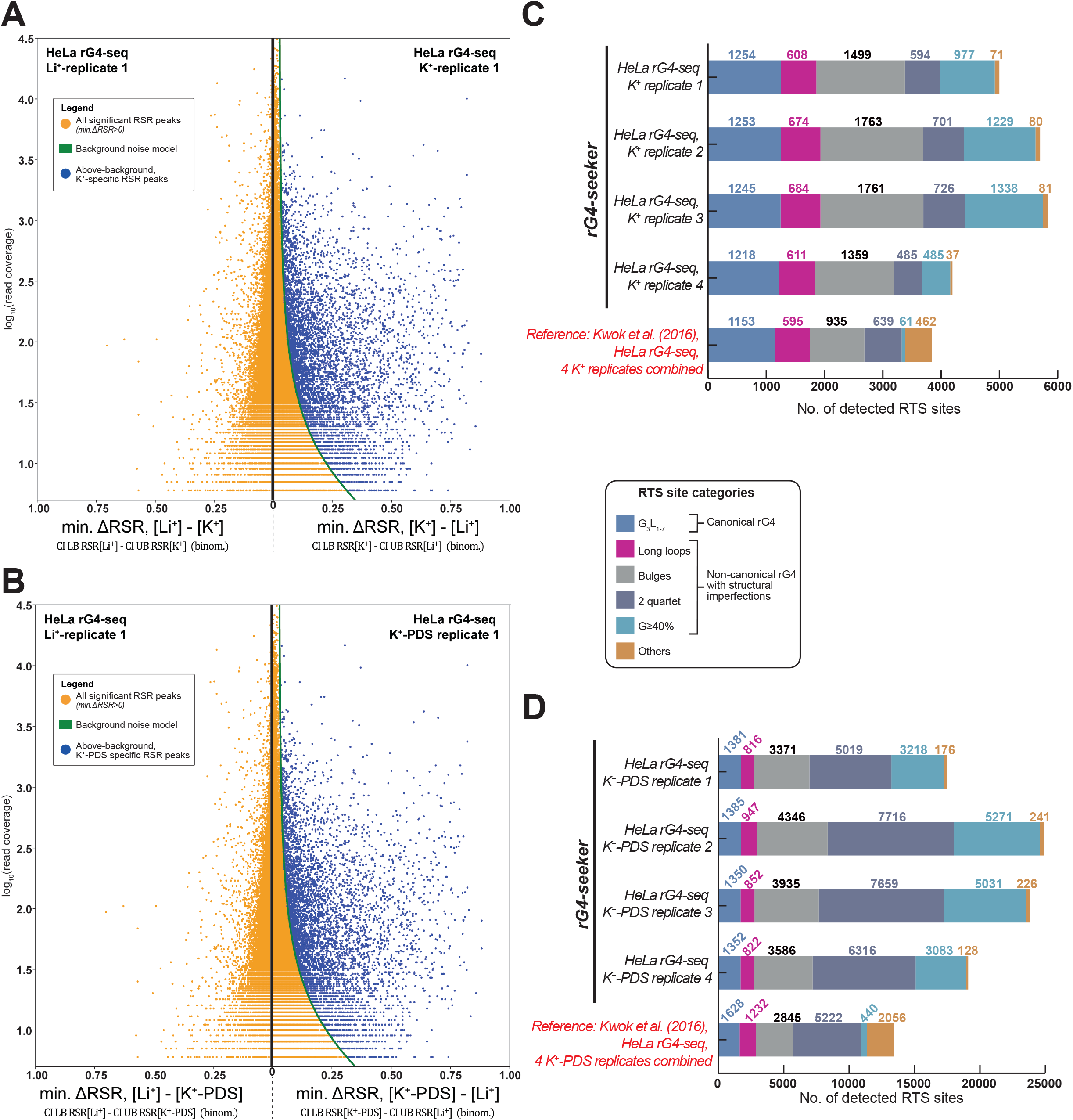
Novel rG4-seq analysis workflow enables reliable, replicate-free RTS site detection. (a, b) RSR peaks identified *ab initio* by applying the minimum ΔRSR metric scheme, sequence-based filtering scheme and fragmentation-associated noise model. (a) RSR peaks detected from replicate 1 of HeLa rG4-seq dataset (pairwise comparison between K^+^/Li^+^ conditions). (b) RSR peaks detected from replicate 1 of HeLa rG4-seq dataset (pairwise comparison between K^+^-PDS/Li^+^ condition). (c, d) Summary of RTS sites detected each replicate rG4-seq dataset, categorized by associated rG4 structural motifs for each site. Adjacent, consecutive RSR peaks were merged and considered to indicate a single RTS site. RTS site detection results (combined analysis of four replicates) from Kwok et al. (2016) were included as a reference. (c) RTS sites detected from pairwise comparison between datasets of K^+^/Li^+^ conditions (d) RTS sites detected from pairwise comparison between datasets of K^+^-PDS/Li^+^ conditions.

The RTS sites were called in a replicate-independent manner under a stricter FDR setting. Compared with our previous analysis combining four replicates, the new workflow detected a similar number of RTS sites associated with the canonical/G3L_1-7_ and long loop rG4 structural motifs and revealed more sites belonging to the bulges, 2-quartet and G≥40% motifs (Figure 4C, D). These findings suggested that the new RTS calling workflow out-performed previous methods by easing the replicate requirements and fully exploiting the potential of rG4-seq technology at any sequencing read coverage.

### rG4-seeker pipeline offers improved integrative analysis for rG4-seq dataset

These findings and proposed workflows were further integrated into a new rG4-seq bioinformatic analysis pipeline called rG4-seeker. Using a genome-aligned rG4-seq dataset as input, rG4-seeker automatically conducted replicate-independent RTS site calling and reported a list of replicate-consensus RTS sites classified by their associated putative rG4 motifs. The re-analysis of published HeLa rG4-seq datasets using rG4-seeker revealed 5,528 consensus RTS sites that agreed between replicate datasets and between K^+^ and K^+^-PDS experimental conditions, with only 14 of these sites assigned to the “Others” class (false-positive detections) (Figure 5).

**Figure 5.**
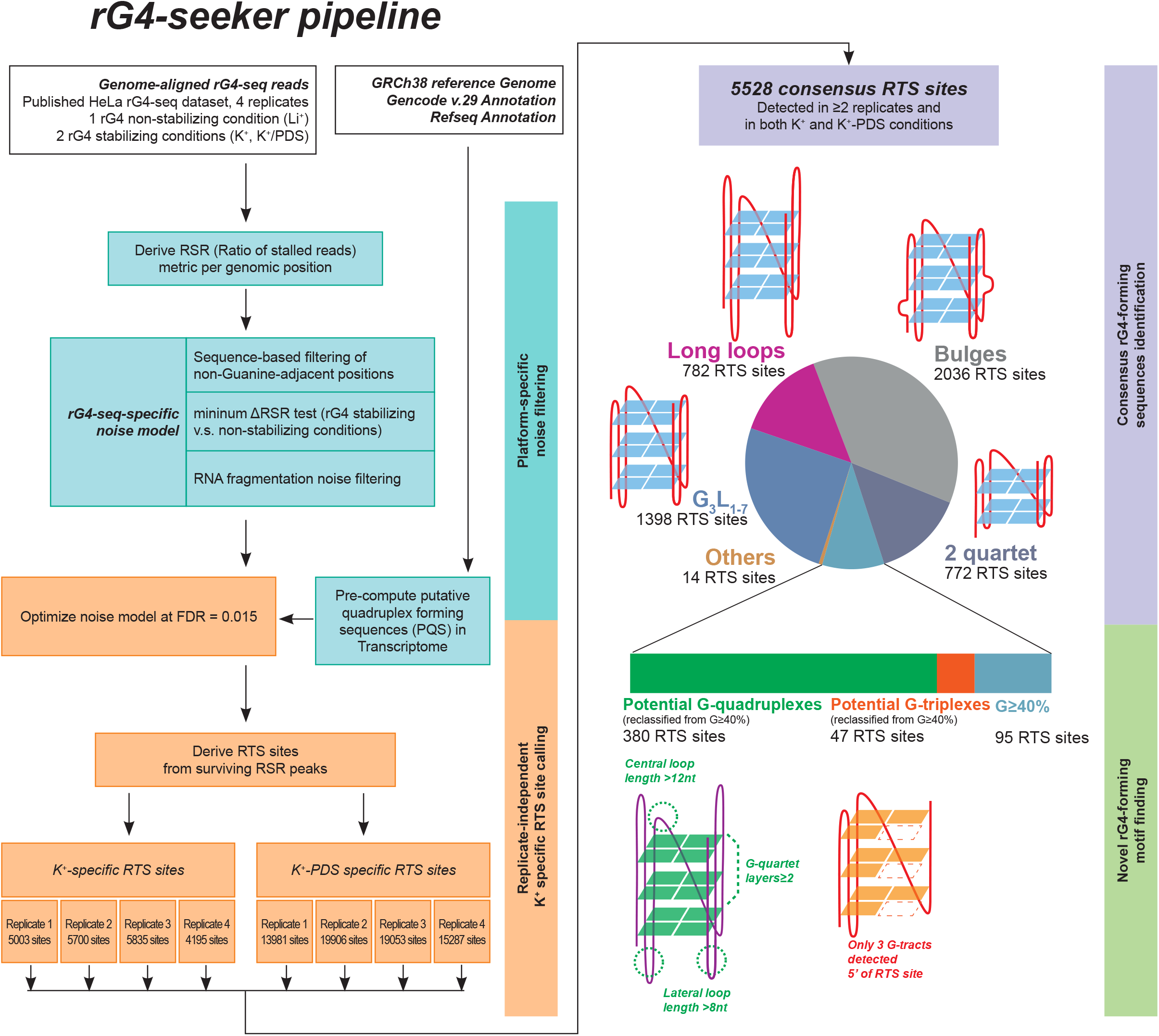
Workflow of rG4-seeker pipeline on published HeLa rG4-seq dataset. rG4-seeker is a novel pipeline for rG4-seq analysis that automates the proposed workflow of RTS site detection (minimum ΔRSR metric scheme, sequence-based filtering scheme and fragmentation-associated noise model) and combines detection results from multiple rG4-seq replicate experiments by consensus.

### Novel patterns of structural imperfection toleration in rG4 motifs

From the HeLa rG4-seq dataset, rG4-seeker identified 380 novel RTS sites with adjacent sequences that resembled rG4s but did not match the canonical/G3L_1-7_ motif and non-canonical motifs (long loops, bulges and 2-quartet). We found that the adjacent sequences of these novel RTS sites could be described using a more lenient motif definition that tolerated multiple structural imperfections simultaneously (Figure 5). Meanwhile, another 47 RTS sites were detected with only three G-tracts detected at the 5’ of RTS sites, suggesting that these sites were associated with potential RNA G-triplex (rG3) structures, which is interesting as only DNA G-triplex structures have been reported previously^13,14^.

Two potential rG3 and four potential rG4 candidates were selected for experimental validation using RTS PAGE assay. All candidates induced RTS at the genomic positions as indicated by rG4-seeker analysis (Figure 6A). The two candidate RTS sites located on *MAGOHB* (Figure 6B) and *ANP32E* (Supplementary Figure S7A) genes were previously considered to be “Others” but were re-classified as potential rG4 and rG3 by rG4-seeker. Interestingly, RTS PAGE assay revealed that both potential rG3s on *DAG1* (Figure 6C) and *PCNP* (Supplementary Figure S7D) genes were *de facto* G-quadruplexes, with their 3’-most G-tracts located 27 nt and 13 nt downstream of the remaining G-tracts that were not discovered by rG4-seq. Meanwhile, for three of the four potential rG4 candidates, gel bands corresponding to RT stalling events were observed adjacent to both the 3^rd^ and 4^th^ (3’-most) G-tracts, suggesting that the observations from the two potential rG3 candidates were a general phenomenon. One possible explanation would be the presence of G-triplex intermediates during G4 folding as suggested by multiple empirical studies^15,16^.

**Figure 6.**
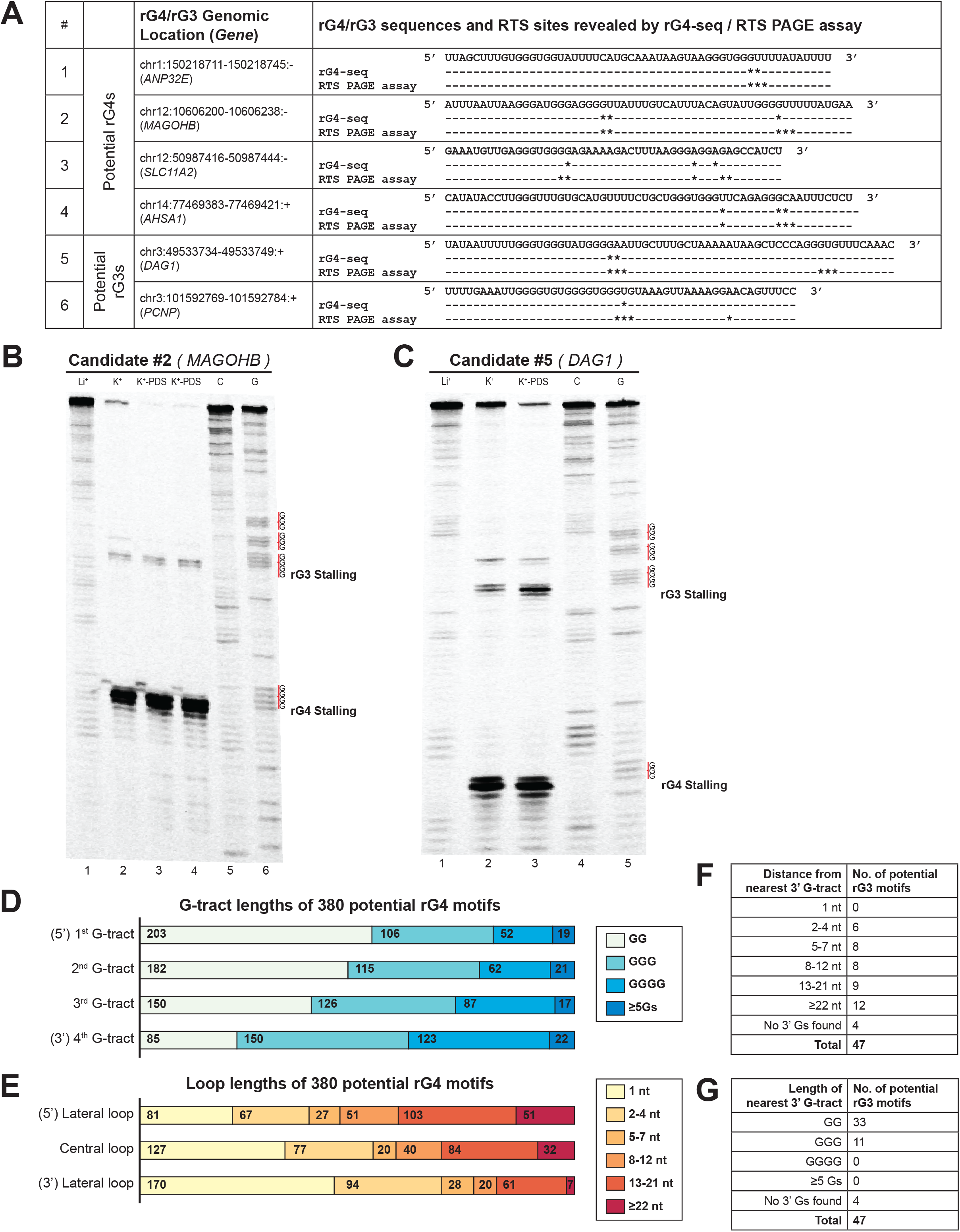
Detected potential G-quadruplexes and G-triplexes suggest novel patterns of structural imperfection toleration in rG4 motifs. (a) Summary of six potential G-quadruplexes (rG4)/G-triplexes (rG3) candidates selected for RTS PAGE validation. Location of RTS sites revealed by rG4-seq and RTS PAGE assay were indicated separately (* represents one nucleotide position where RTS is detected). Most RTS sites discovered by rG4-seq were reproduced in the RTS PAGE assay. (b) RTS PAGE assay of a potential rG4 candidate on *MAGOHB* gene indicated clear RTS at the 3’ end of the 3^rd^ and 4^th^ G-tracts, respectively. (c) A potential rG3 was initially suggested to form on *DAG1* gene based on findings from rG4-seq. RTS PAGE assay revealed a 4^th^ G-tract 27 nt downstream of the 3^rd^ G-tract, indicating that the region harbored a rG4 motif instead. (d, e) Summary of G-tract and loop lengths of 380 potential G-quadruplexes. The results suggest that toleration for structural imperfection (less than 3 Gs in G-tracts, and longer loops) was higher in the 5’ region and lower at the 3’ end of rG4 motifs. (f, g) Summary of 3’ downstream G-tracts of 47 potential G-triplexes identified by nucleotide sequence searching. 3’ downstream G-tracts were identified in most (43 out of 47) of the rG3s, where most G-tracts were composed of 2 Gs.

To better understand the structural imperfection tolerance among the potential rG4s identified, we summarized and compared the G-tract lengths and loop lengths of these 380 motifs, and revealed associations between their location in a rG4 motif and their respective lengths (Figure 6D, E). We found that while the length of a G-tract positively correlated to its distance from the 3’ end of the motif, the length of the connecting loop negatively correlated to its distance from the 3’ end of the motif. Moreover, we found that most of the 47 potential rG3s had a 4^th^ G-tract at the 3’ end of the RTS site, with most of the G-tracts composed of two Gs (Figure 6F, G). These observations suggested that the structural imperfections at the 3’ side of rG4s prevented detection by rG4-seq, and that the 3’ side of rG4s is less tolerant of structural imperfections.

### Reproducibility of rG4-seq experiments affected by stochasticity of rG4-induced RTS effects

The replicate-independent analysis workflow in rG4-seeker enabled us to investigate the reproducibility of RTS site detection in the rG4-seq experiments. By surveying the detection status of 5,528 report RTS sites among the eight rG4-seq datasets (four replicates in K^+^ and K^+^-PDS conditions), we found that only ~2,000 RTS sites were detected unanimously among all datasets (Figure 7A). Increasing the sequencing read coverage only partially improved the reproducibility of RTS site detection: while reproducibility was greatly improved by coverage increments between 6x and 32x, the marginal returns for increments beyond 32x greatly diminished (Figure 7B). Notably, at read coverages of 512x to 1,024x or above, 40% of RTS sites were not reproduced in at least one of the four replicates at K^+^ conditions. These observations indicated a random uncertainty in whether an rG4 motif would reproduce RTS in an rG4-seq experiment, despite all experimental conditions being kept constant, suggesting stochasticity in rG4s’ induction of RTS.

**Figure 7.**
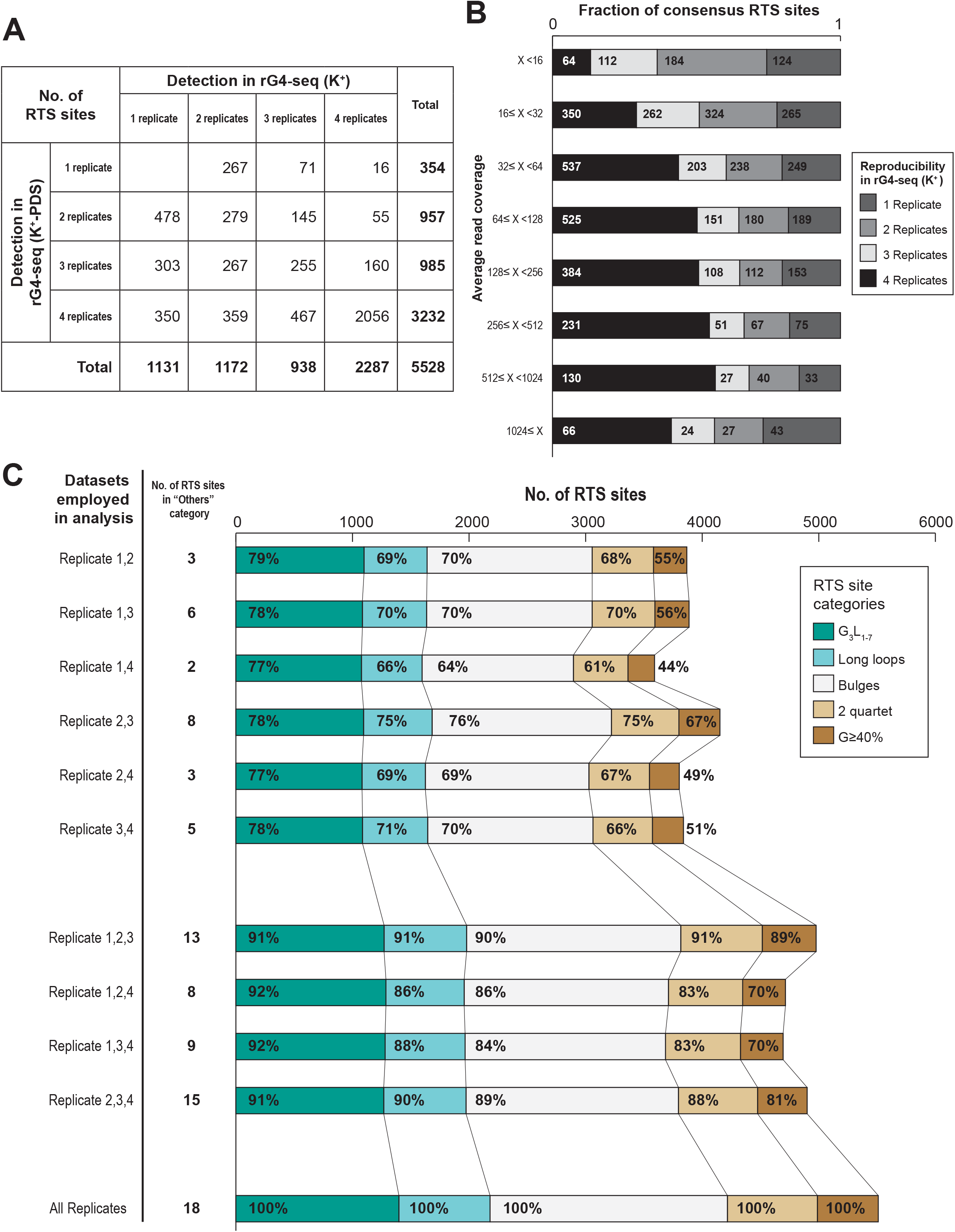
rG4 induces RTS effect in stochastic manner and contributes to divergence between replicates. (a) Breakdown of 5528 detected consensus RTS sites by their reproducibility in K^+^ and K^+^-PDS replicate datasets. Only ~40% of RTS sites were simultaneously detected in all four replicates and in both K^+^ and K^+^-PDS conditions. (b) Distribution of 5528 consensus RTS sites by average read coverage (logarithmic scale) and detection status at the K^+^ condition. While RTS sites of higher read coverage in general exhibited higher reproducibility in rG4-seq (K^+^), the marginal improvement in reproducibility by increasing read coverage beyond 32x was subtle. (c) Comparison of rG4-seeker analysis results using only a subset of two or three replicate datasets among four available replicates. Reduced number of replicates caused fewer RTS sites detection but did not increase the detection of RTS sites in the “Others” category (false-positive detections).

To better illustrate the stochasticity of rG4-induced RTS, we repeated rG4-seeker analysis using combinations of only two or three replicate HeLa rG4-seq datasets (Figure 7C). We found that the lower number of replicates reduced the total number of identified rG4-induced RTSs: compared to a four-replicate analysis, a two-replicate analysis only reproduced ~70% of all RTS sites, while a three-replicate analysis reproduced up to 90% of all RTS sites. Moreover, the structural motifs of the rG4s also appeared to influence reproducibility when fewer replicates were used: the reproducibility was in general higher for canonical/G3L_1-7_ motifs but lower for those with structural imperfections, indicating that the stochasticity of induced RTS depended on the stability of the rG4s. Taken together, the results suggested that a two-replicate rG4-seq experiment was sufficiently reliable for transcriptome-wide rG4 mapping, and that the use of more replicates improved the sensitivity for rG4s that induced RTS in a stochastic manner. Lastly, genuine rG4s were more likely to be reproducibly detected by rG4-seq if the read coverage was 32x or more.

### High local GC% may compromise RTS site detection by rG4-seq

The RTS calling results from rG4-seeker and prior methods were compared to derive the recalling statistics for RTS sites. We found that among the 1,143 and 2,215 RTS sites (K^+^) previously reported to be associated with canonical/non-canonical rG4 motifs, approximately 20–25% of them were not recalled in the rG4-seeker re-analysis (Figure 8A). Interestingly, the adjacent sequences of the non-recalled RTS sites appeared to have substantially higher GC% than the recalled RTS sites (Figure 8B): in both the 5’ upstream and 3’ downstream regions, the average GC% for non-recalled RTS sites fell to around 60% compared to that of 50-55% for recalled RTS sites. We speculated that this was a result of the GC-bias behavior of Illumina sequencing technology, where GC-rich fragments are typically underrepresented^17,18^. Thus, given that the prior arts utilized local coverage drops to identify RTS events, it is likely that the algorithm overlooked GC-bias and RTS-induced coverage changes. In contrast, rG4-seeker was not affected by GC-bias. To accurately detect rG4s that reside within high GC regions, RTS PAGE analysis or alternative sequencing technologies are required.

**Figure 8.**
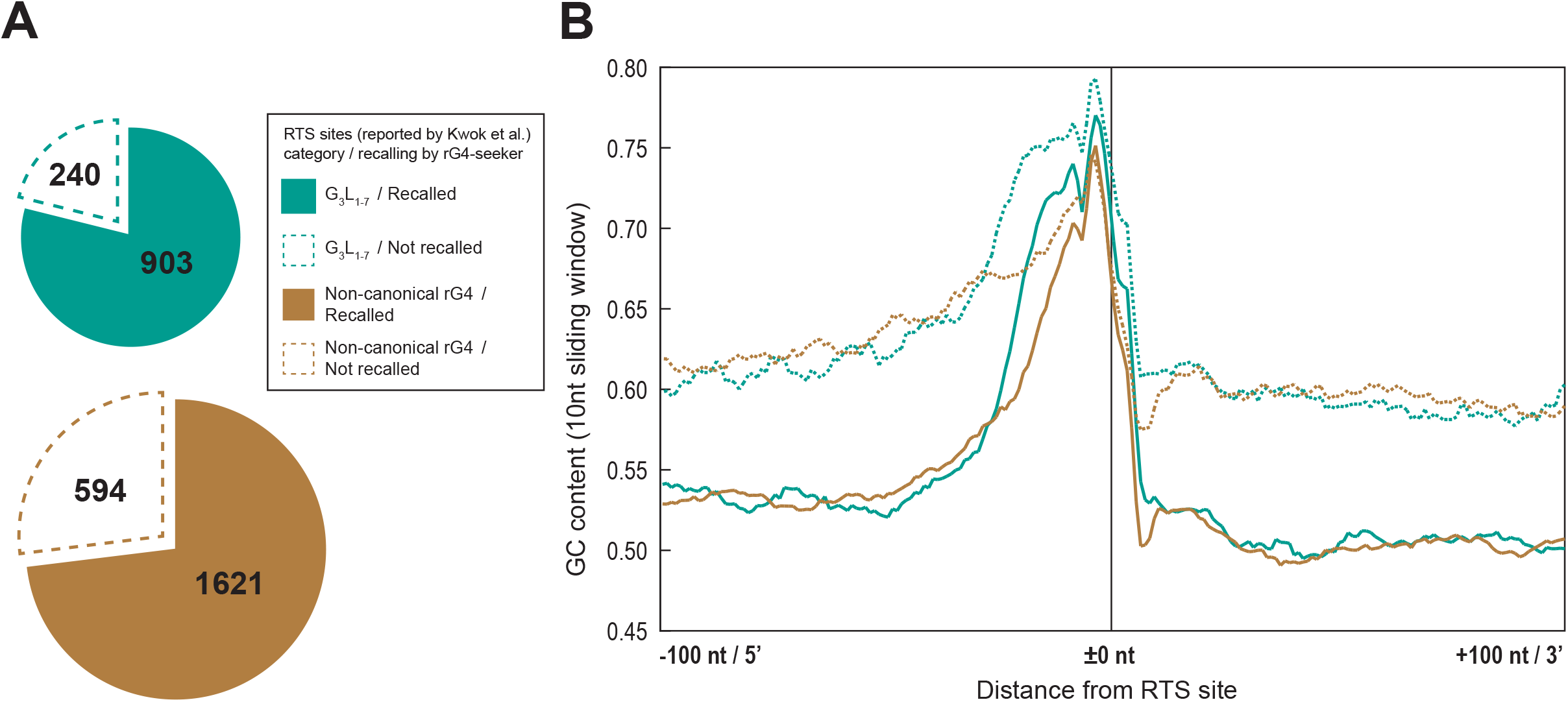
High local GC% may compromise RTS site detection with rG4-seq. (a) Recall statistics of RTS sites (K^+^) from canonical and non-canonical rG4 categories reported by Kwok et al. (2016). Around 20–25% of RTS sites were not recalled by rG4-seeker in the re-analysis. (b) Average GC% (10 nt sliding windows) in the +-100 bp region of 3358 RTS sites reported by Kwok et al. (2016), segregated based on their motif categories and recall status by rG4-seeker. RTS sites that were not recalled had higher GC% across the entire region compared with recalled RTS sites.

## DISCUSSION

We developed rG4-seq technology to address the limitations of low-throughput rG4 experimental assays and rG4 computational prediction algorithms. It offers empirical high-throughput detection and mapping of *in vitro* rG4 structures on a transcriptome-wide scale. The rG4-seeker pipeline was engineered with two objectives in mind: to enable associative analysis to infer *in vivo* rG4 biological functions, and to offer a screening for high confidence rG4 candidates for functional studies.

In terms of the first objective, the noise model of rG4-seeker was designed to be conservative in detecting rG4s of lower RTS strength as a compromise for a lower false-positive detection rate. The tradeoff would offer better promise for downstream applications, such as machine learning applications to extrapolate rules governing rG4 formation, or associate studies that connect rG4s with other RNA regulatory and sequence elements. Under this conservative scheme, the rG4-seeker made >2,000 novel rG4 detections. Interestingly, most of these newly identified rG4s were non-canonical motifs, had structural imperfections, and were expected to be less thermodynamically stable. The greater prevalence of non-canonical rG4s in the transcriptome compared with canonical rG4s agreed with the findings on selection pressures against stable genomic and transcriptomic G4 sequences^19,20^. Moreover, as the free energy differences between folded and unfolded conformations of less stable rG4s were typically lower, they might be more relevant in processes that involve dynamic folding/unfolding of rG4 structures than more stable motifs.

In terms of the second objective, the replicate-independent nature of rG4-seeker would significantly reduce the cost of rG4s screening within rG4-seq dataset. Our results indicate that a two-replicate rG4-seq experiment is sufficient to eliminate noise, and that the K^+^-PDS condition was not required. These factors implied that rG4-seq experiments could be down-scaled from 12 (four replicates, three conditions) to 6 (two replicates, three conditions) or 4 (two replicates, two conditions). For rG4-seq validation of existing putative quadruplex sequences-of-interest, rG4-seeker can allow visualization of RSR signals in a genome browser analogous to a RTS PAGE assay gel image and show the RTS information of all adjacent positions near any PQS-of-interest (Supplementary file 2).

Finally, our analysis revealed that the sampling error and background noise observed in rG4-seq were due to RNA-seq library preparation procedures but not the reverse transcription buffer/ionic conditions. The error and noise were consequences of sequencing read coverage variations owing to differences in transcript abundances, RNA fragmentation procedure in library preparation, and the sequence coverage bias of Illumina technology. These issues may in particular affect high-throughput RNA sequence/structure probing technologies that use RTS as readouts and require bioinformatic analysis at single-nucleotide resolution. Our study demonstrates that simple statistical models based on binomial distributions are sufficient to address these issues without additional control experiments. Our method can also be incorporated into the bioinformatic pipeline of other RNA-seq technologies to improve analysis outcomes.

## CONCLUSIONS

The rG4-seq method was developed to detect *in vitro* RNA G-quadruplex (rG4s) structures on a transcriptome-wide scale. However, the application of rG4-seq has been confined by its requirements of high number of replicates and a moderate false-positive rate. To alleviate these limitations and to improve the outcomes of rG4-seq experiments, we examined rG4-seq chemistry and established models for both indigenous sampling errors and background noise in rG4-seq data. We developed rG4-seeker, a novel bioinformatics pipeline for rG4-seq analysis. Compared with previous methods, rG4-seeker exhibited better false-positive discrimination and improved sensitivity for rG4s calling. This method is also less dependent on biological replicates. Using rG4-seeker, we identified novel features in the HeLa rG4-seq dataset that were previously missed, and experimentally validated selected cases. rG4-seeker empowers reliable rG4-seq investigation at a reduced cost, laying the foundations for future development of rG4 biology.

## MATERIALS AND METHODS

### Bioinformatic preprocessing of rG4-seq data

The HeLa rG4-seq dataset generated previously^9^ was used in this study. Sequencing reads were first adapter-trimmed and quality-trimmed using cutadapt^21^. Trimmed reads were then aligned to GRCh37/hg19 or GRCh38/hg38 human reference genome using STAR aligner^22^, and only uniquely aligned reads were used for subsequent analysis. Read coverage and number of read starts per nucleotide position were counted using Python. RefSeq^23^ and GENCODE v.29 ^24^ gene annotation databases were used to define the transcriptomic regions.

### Defining the ratio of stalled reads (RSR) metric

The RSR measurement at each sequenced nucleotide position was calculated using the following formula:

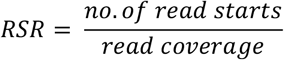

The RT stalling/read-through event at a position can be modelled as a Bernoulli process, where

- probability *p* indicates the underlying chance of RT stalling at the position, and is estimated by RSR measured from rG4-seq experiments,
- each sequencing read covering the position corresponds to an independent Bernoulli trial, and
- the trial is a “success” if the read start is also located at the position, otherwise it is a “failure.” Read start counts observed in a position can thus be assumed to follow a binomial distribution B(*n*, *p*), where *n* is the read coverage and *p* is the underlying probability of RTS events.

### Binomial test with RSR metric

As read start counts observed in a single nucleotide position will follow a binomial distribution, a one-sided binomial test was applied to compare RSR measurements between rG4-stablizing (K^+^ or K^+^-PDS) and rG4-non-stabilizing (Li^+^) conditions for the following hypotheses.

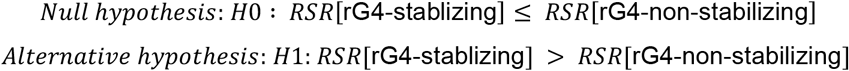

A statistically greater RSR measurement at rG4-stablizing conditions suggests that the nucleotide position corresponds to an rG4-induced RTS event, which is expected to occur at the 3’ end of the rG4. In contrast, when the RSR measurement is not significantly increased by rG4-stablizing conditions, this suggests that the nucleotide position is either in a transcript region that does not support rG4 formation or located within an rG4 motif but away from the 3’ end.

### Reads subsampling

Random read subsampling was conducted using samtools^25^ on highly expressed gene regions of read coverage >10000x.

### Minimum ΔRSR scheme

Given that the RSR measurement at any read coverage follows a binomial distribution, binomial proportional confidence intervals (CIs) were applied to estimate the minimum RSR at rG4-stablizing (K^+^ or K^+^-PDS) conditions (using the lower bound value) and the maximum RSR at rG4-non-stabilizing (Li^+^) conditions (using the upper bound value). If the difference between the two values (named “minimum ΔRSR”) is greater than zero, then RSR[rG4-stabilzing] > RSR[rG4-non-stabilizing], and the RSR peak is statistically significant. Essentially, the scheme is equivalent to substituting *p* used in the binomial test with its upper-bound estimation from the binomial-proportional CI, thus being more conservative at low read coverage.

### Re-analysis of RTS sites in “Others” category

We previously assigned 462 RTS sites (K^+^) and 2056 RTS sites (K^+^-PDS) to the “Others” structural subclass category. Based on the context of adjacent sequences, the majority of these “Others” RTS sites were predicted to be false positives rather than genuine rG4 structures^9^. Re-analysis of these RTS sites using the RSR metric and binomial test/minimum ΔRSR scheme was conducted to evaluate the efficacy of the new detection scheme in rejecting these false-positive RTS sites.

To match the previous analysis workflow, rG4-seq reads uniquely aligned to the GRCh37/hg19 reference genome were used in the re-analysis. Read coverage and read start counts in a ±2 nt window at the reported “Others” RTS sites (5 nucleotide positions in total) were extracted from each replicate HeLa rG4-seq dataset. Positions of read coverage >6x were considered. Each nucleotide position was then individually tested for statistically significant RSR peaks (where RSR[rG4-stablizing] > RSR[rG4-non-stabilizing]) that passed either the binomial test (one-sided, *p* <0.05) or the minimum ΔRSR scheme (minimum ΔRSR >0, α = 0.1). If at least one associated significant RSR peak was detected at or 3’-adjacent to a guanine residue, the parent RTS site was considered adjacent to that guanine residue. Given that PDS is a small molecule with a different G4-stabilizing mechanism than cations^26^, we reasoned that if a genuine rG4 can fold in K^+^ condition, then it should also fold in K^+^-PDS condition. Considering this property, the 462 “Others” RTS sites (K^+^) and 2056 “Others” RTS sites (K^+^-PDS) were overlapped. Our results revealed 42 “others” RTS sites that were detected in both conditions. Within these 42 detected RTS sites, 35 were located at or adjacent to guanine residues (Supplementary Table S1), among which 24 sites were adjacent to 1 or more GGG motifs that resembled G-tracts (Supplementary Table S1). Furthermore, 2 sites on *ANP32E* and *MAGOHB* genes were later verified with an RTS PAGE assay to correspond to genuine rG4s (see results). These findings suggested that some of these 35 simultaneously detected “Others” RTS sites were rG4s misclassified as false positives. In contrast, the remaining 7 simultaneously detected RTS sites were not adjacent to any guanine residues, indicating that they were genuine false positives.

In the subsequent re-analysis using the RSR metric and binomial test/minimum ΔRSR scheme, we explicitly overlapped the RTS sites detected in K^+^ and K^+^-PDS conditions to evaluate their performance (see results). An ideal RTS site detection scheme should selectively retain some/all of the 29 simultaneously detected “Others” RTS sites associated with rG4s, while rejecting all of the remaining “Others” RTS sites.

### Sequence-based filtering scheme

Based on the observation that rG4-induced RTS sites occurred only at or 3’-adjacent to a G-tract, a sequence-based filtering scheme was derived to whitelist nucleotide positions that fulfilled the following requirements for RSR peak calling as follows:

1. At least 1 G residue at −2 nt, −1 nt or 0 nt positions
2. At least 3 G residues or 1 “GG” motif between −9 nt and 0 nt positions

The criteria limited RSR peak calling to positions at or with 2 nt 3’ of a G-tract, where a G-tract could be either composed of ≥2 consecutive Gs (corresponding to the canonical/G3L_1-7_, long loops and 2-quartet motifs), or ≥3 Gs but interrupted by a bulge of 1 to 7 nts (corresponding to the bulges motif). The whitelisting of nucleotide positions was limited to annotated, exonic transcriptomic regions and repeated for each annotated transcript isoform. The sequence of the 5’ adjacent exon was considered if the surveyed nucleotide position was adjacent to an annotated splice junction. The union was taken to be between individual whitelisting results from overlapping genes/isoforms.

### Statistical modelling of fragmentation-associated background noise

The probabilistic upper limit of fragmentation-associated background noise was modelled as follows.

1. Estimate the average expected RSR measurement for any sequenced nucleotide position with the conditional probability *P* (*A* | *B*), where

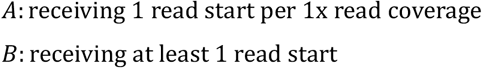 The conditional probability *P* (*A* | *B*) was further derived from two probabilities using Bayes’s theorem (Supplementary Table S2):

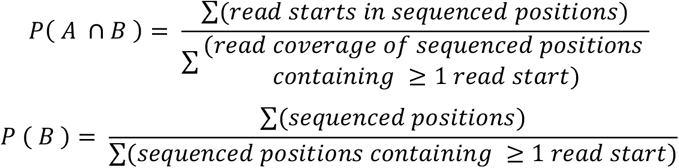 Although the respective contribution of RT stalling and fragmentation noise to the RSR is not understood, a worst-case scenario in which it is solely contributed by fragmentation noise can be assumed.
2. Estimate the maximum possible RSR measurement contributed by fragmentation noise using the upper bound value of confidence interval of binomial distribution *B*(*n* = *read coverage*, *p* = *P* (*A* | *B*)). Similar to RT stalling events, RT completion events at the 5’ end of RNA fragments at any nucleotide position can also be modelled as a Bernoulli process.
3. Calculate the maximum difference in RSR owing to fragmentation-associated noise (max. ΔRSR[fragmentation]) between two libraries, which corresponds to the probabilistic upper limit. The max. ΔRSR[fragmentation] is equal to the upper CI bound of *B*(*n* = = *read coverage*, *p* = *p* = *P*(*A* | *B*)) as derived in 2, because in the worst-case scenario, one library would receive maximum fragmentation-associated noise while the other would receive zero noise.

To estimate *P* (*A* | *B*), the read coverage and distribution of read starts across the entire sequenced transcriptome were measured and summarized. We found that when considering only regions with read coverage ≥2048x, around 95% of surveyed nucleotide positions received 1 or more read starts, where the conditional probability *P* (*A* | *B*) fell between 1.18% to 1.44%, and were close to the theoretical *P* (*A* | *B*) of 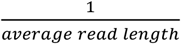 (Supplementary Table S2, S3). However, when all sequenced regions (read coverage ≥6x) were considered, only around 30–36% of positions were found to receive 1 or more read starts, where *P* (*A* | *B*) fell between 4.42% and 5.70% (Supplementary Table S2). As most of the RSR peaks with high minimum ΔRSR values were found at regions of coverage between 6x and 2048x, their geometric mean of approximately 2.5% was chosen as the global *P* (*A* | *B*) for deriving the upper limit of fragmentation noise across all libraries.

### *Ab initio* replicate independent RSR peak identification and RTS site calling

Pairwise comparisons between K^+^/Li^+^ and K^+^-PDS/Li^+^ dataset in each HeLa rG4-seq replicate were conducted by applying the minimum ΔRSR metric scheme, sequence-based filtering scheme and fragmentation-associated noise model as described above. Statistically significant RSR peaks (RSR[rG4-stabilizing] > RSR[rG4-non-stabilizing]) were successively derived, and consecutive adjacent RSR peaks merged. After merging, each RSR peak was considered an RTS site that spanned 1 or more nucleotide positions. Detected RTS sites were then classified into 6 categories (canonical/G3L_1-7_, long loops, 2-quartet, bulges, G≥40% and Others) based on their adjacent sequences and according to the scheme described in our previous work^9^.

The α values for deriving the binomial proportional confidence intervals (CIs) for minimum ΔRSR scheme and fragmentation-associated noise model were automatically optimized for a target FDR level (assuming RTS sites classified as “Others” to be false positives) by the following logic:

1. The minimum ΔRSR test would be repeated using 7 preset α values (0.2, 0.159, 0.126, 0.1, 0.079, 0.063, 0.05) that ranged between 0.2 and 0.05.
2. For each trial of minimum ΔRSR test, the optimal *α* value for the fragmentation-associated noise model would be derived by exhaustion. The *α* value that satisfied the target FDR level and enabled detection of the greatest number of RTS sites was considered optimal.
3. Among the 7 trials of minimum ΔRSR test, the trial and the 2 α values that enabled detection of the greatest number of RTS sites were considered the global optimal results.

### rG4-seeker pipeline

The rG4-seeker pipeline was implemented in Python, which integrated and automated all of the procedures of rG4-seq analysis described in this study, including

1. generation of transcriptome-wide sequence-based filtering scheme,
2. pre-computation of putative quadruplex forming sequences (PQSs) according to rG4 structural motifs (canonical/G3L_1-7_, long loop, bulges and 2-quartet), and
3. replicate-independent RSR peak identification/RTS site calling workflow. Using the rG4-seeker pipeline, we generated a list of consensus RTS sites that were detected in ≥rG4-seq replicates and in both K^+^ and K^+^-PDS conditions.

### Re-classification of G≥40% RTS sites

All RTS sites in the G≥40% category were tested for potential re-classification as G-quadruplexes or G-triplexes based on the 50 nt sequence 5’ of the RTS site as follows:

1. If 4 G-tracts of ≥2 consecutive Gs were found within the 50 nt sequence, and the RTS site was within 2 nt of the 3’-most G-tract, the RTS site was reclassified into the “potential G-quadruplex” category; otherwise,
2. If 3 G-tracts of ≥2 consecutive Gs were found with the 50 nt sequence, and the RTS site was within 2nt of the 3’-most G-tract, the RTS site was reclassified into the “potential G-triplex” category.

### Preparation of DNA oligonucleotides for spectroscopy and luciferase assays

DNA oligonucleotides for RNA preparation and RTS assay were purchased from Integrated DNA Technologies, USA and Genewiz, China.

### Preparation of *in vitro* transcribed (IVT) RNAs

DNA hemi-duplex for in vitro transcription purpose was produced after hybridization of 20 uM T7 DNA promoter and 20 uM reverse DNA template. IVT RNAs was generated using DNA hemi-duplex and a HiScribe T7 High Yield RNA Synthesis Kit following the protocol described online (New England Biolabs, USA). RNA was purified using precast 15% denaturing polyacrylamide gel (Thermo, USA). The desired RNA band was cut under brief UV shadowing, and crush and soaked in 1× 10 mM Tris-HCl pH 7.5, 1 mM EDTA, 800 mM LiCl (1X TEL800) in a thermoshaker at 4 °C, 1300 rpm overnight. The gel piece were filtered by microcentrifuge filter, and the RNA was then extracted and purified using RNA Clean & Concentrator-5 following the protocol described according to the manufacturer’s protocol (Zymo Research, USA). The RNA is quantified by nanodrop and stored in −20°C before use.

### PAGE-based RTS assay

A 10 uL reaction including 5 pmol IVT RNA, 5 μM Cy5-labelled DNA primer and reverse transcription buffer (150 mM KCl, 4 mM MgCl_2_, 20 mM Tris pH 7.5, 1 mM DTT, and 0.5 mM dNTPs) were mixed and heated to 75 °C for 3 min and then to 35 °C for 5 min. The reaction was equilibrated at 37°C for 10 min, then 0.5 μL of Superscript III (200 U/μL) (Thermo Fisher Scientific, USA) was added and incubated at 50 °C for 15 min before 0.5 μL of 2 M NaOH was added. After heating at 95 °C for 10 min, 2X denaturing formamide dye was added to stop the reaction. cDNAs were size fractionated by 8 M urea 8% polyacrylamide gel, and subsequently imaged using Fujifilm FLA-9000.

## AVAILABILITY

rG4-seeker is open source software available in the GitHub repository (https://github.com/TF-Chan-Lab/rG4-seeker)

## SUPPLEMENTARY DATA

Supplementary File 1: Supplementary Figure S1-7 and Supplementary Table S1-3

Supplementary File 2: List of 5,528 RTS sites detected from HeLa rG4-seq dataset by rG4-seeker (Microsoft Excel Spreadsheet)

## ACKNOWLEDGMENTS

Not applicable.

## AUTHOR CONTRIBUTIONS

EYCC, CKK and TFC had the initial idea for the project. EYCC designed and implemented the rG4-seeker pipeline and performed the bioinformatic analysis. TFC contributed to the design of the pipeline. KL and CKK performed the experiments. TFC and CKK supervised the study. EYCC, KL, CKK and TFC wrote the manuscript.

