## Supplementary File 1 for "rG4-seeker enables high-confidence identification of novel and non-canonical rG4 motifs from rG4-seq experiments"

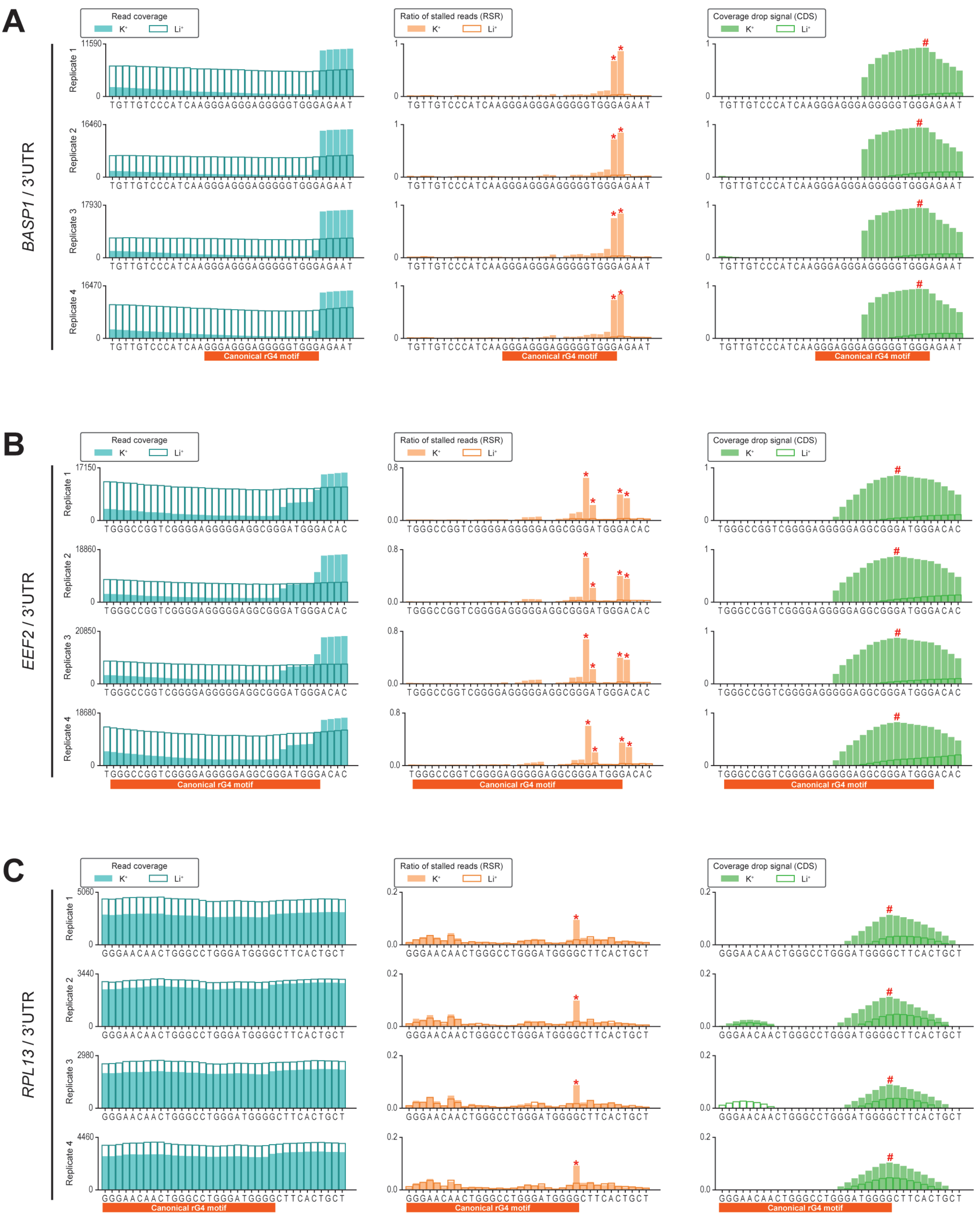

**Figure S1**

Showcases comparing proposed ratio of stalled reads (RSR) metric with coverage drop signal (CDS) metric used to resolve RNA G-quadruplex (rG4)-induced reverse transcriptase stalling (RTS) events at (a) *BASP1* gene (b) *EEF2* gene (c) *RPL13* gene

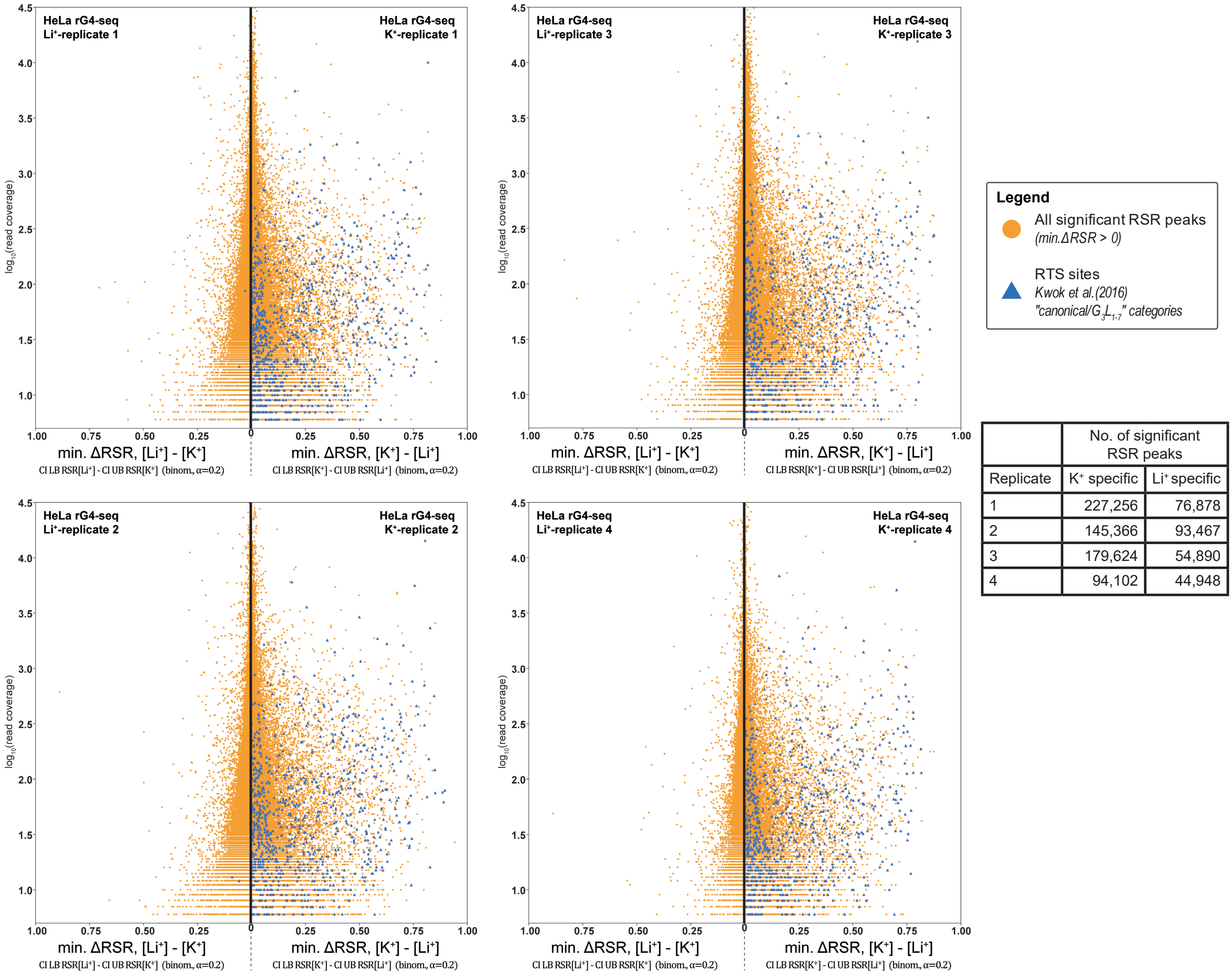

| | No. of significant RSR peaks | |
|---|---|---|
| Replicate | $K^+$ specific | $Li^+$ specific |
| 1 | 227,256 | 76,878 |
| 2 | 145,366 | 93,467 |
| 3 | 179,624 | 54,890 |
| 4 | 94,102 | 44,948 |

**Figure S2**

Transcriptome-wide significant RSR peaks (minimum ΔRSR >0) detected from the 4 replicates of HeLa rG4-seq dataset (pairwise comparison between $K^+$/$Li^+$ condition). Minimum ΔRSR values (indicating RTS effect strength) of RSR peaks were plotted against read coverage (logarithmic scale). Each datapoint corresponds to 1 RSR peak at 1 single-nucleotide genomic locus. RSR peaks coinciding with RTS sites in canonical/$G_3L_{1\text{-}7}$ reported by Kwok et al. (2016) are highlighted.

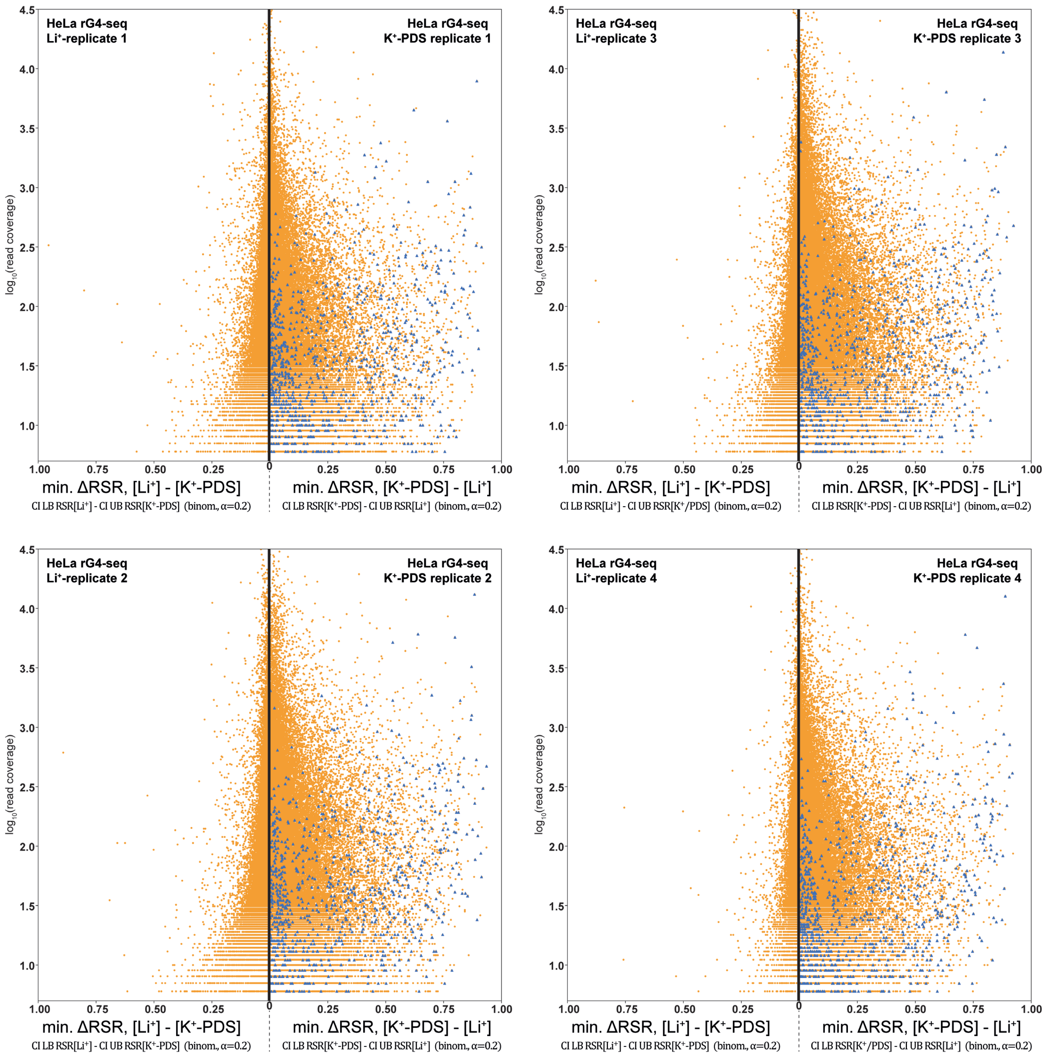

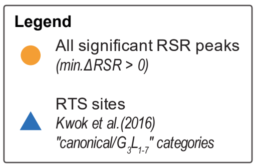

| | No. of significant RSR peaks | |
|---|---|---|
| Replicate | $K^+$/PDS specific | $Li^+$ specific |
| 1 | 285,266 | 89,804 |
| 2 | 181,273 | 128,341 |
| 3 | 199,597 | 77,361 |
| 4 | 155,713 | 35,851 |

**Figure S3**
Transcriptome-wide significant RSR peaks (minimum ΔRSR >0) detected from the 4 replicates of HeLa rG4-seq dataset (pairwise comparison between $K^+$-PDS/$Li^+$ condition). Minimum ΔRSR values (indicating RTS effect strength) of RSR peaks were plotted against read coverage (logarithmic scale). Each datapoint corresponds to 1 RSR peak at 1 single-nucleotide genomic locus. RSR peaks coinciding with RTS sites in canonical/$G_3L_{1-7}$ reported by Kwok et al. (2016) are highlighted.

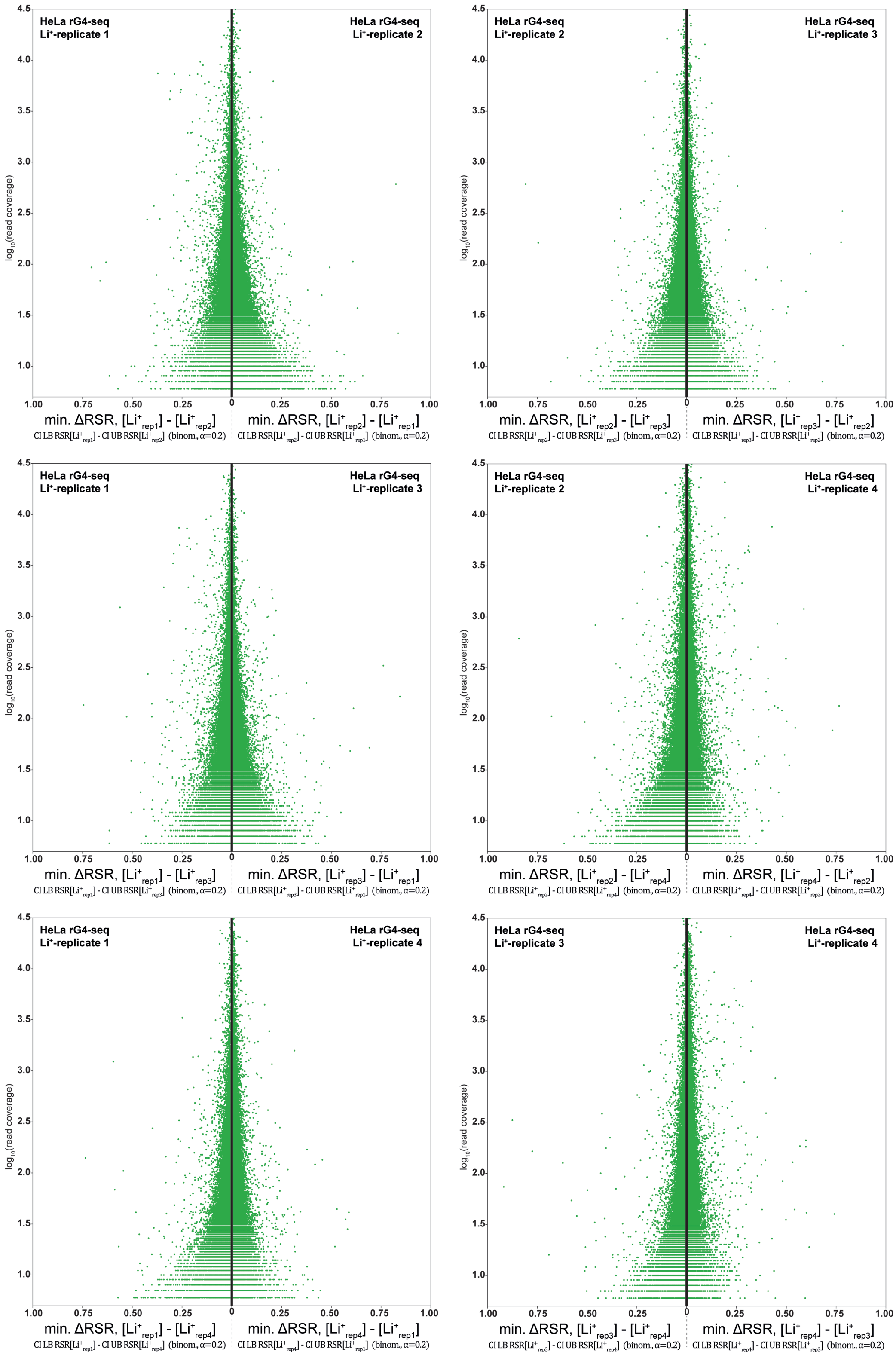

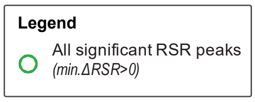

| Pair of Li+ Replicate | No. of significant RSR peaks |
|---|---|
| 1 / 2 | 200,769 / 109,143 |
| 1 / 3 | 149,567 / 110,692 |
| 1 / 4 | 114,978 / 134,213 |
| 2 / 3 | 93,769 / 132,866 |
| 2 / 4 | 78,870 / 161,007 |
| 3 / 4 | 75,363 / 115,232 |

### Figure S4

Transcriptome-wide significant RSR peaks (minimum ΔRSR >0) detected from pairwise comparisons between 4 replicates the $Li^+$ HeLa rG4-seq dataset. Comparison between replicates of identical conditions implies that all detected RSR peaks originated from experimental variations.

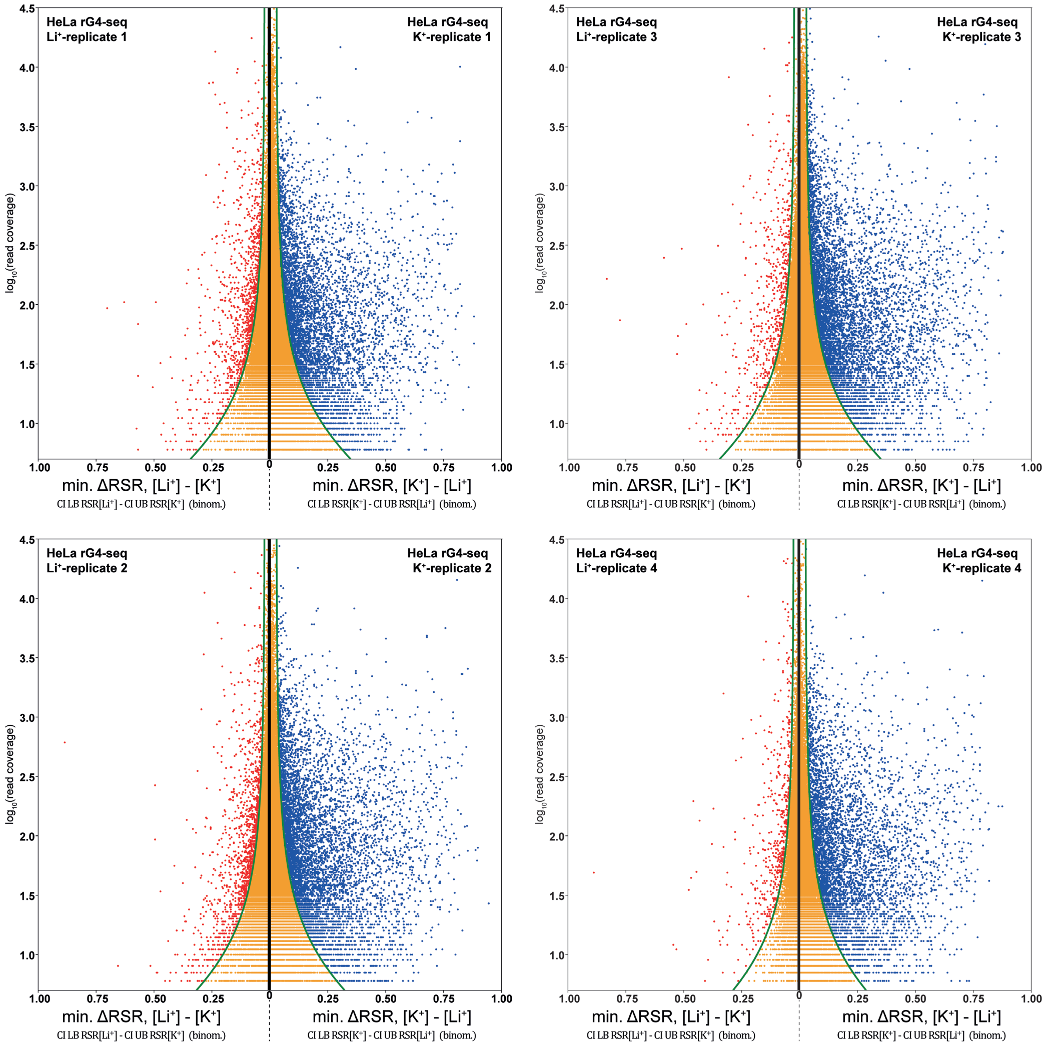

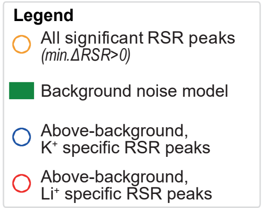

| | Alpha values for binomial proprotion confidence intervals | |
|---|---|---|
| Replicate | min. ΔRSR | Noise model |
| 1 | 0.200 | 0.132 |
| 2 | 0.200 | 0.158 |
| 3 | 0.200 | 0.130 |
| 4 | 0.200 | 0.195 |

| | No. of above-background RSR peaks | |
|---|---|---|
| Replicate | $K^+$ specific | $Li^+$ specific |
| 1 | 5,851 | 1,162 |
| 2 | 6,901 | 1,452 |
| 3 | 6,960 | 701 |
| 4 | 5,195 | 569 |

**Figure S5**

RSR peaks identified ab *initio* from pairwise comparison of 4 replicates of HeLa rG4-seq ($K^+$) and HeLa rG4-seq($Li^+$) by applying the minimum ΔRSR metric scheme, sequence-based filtering scheme and fragmentation-associated noise model.

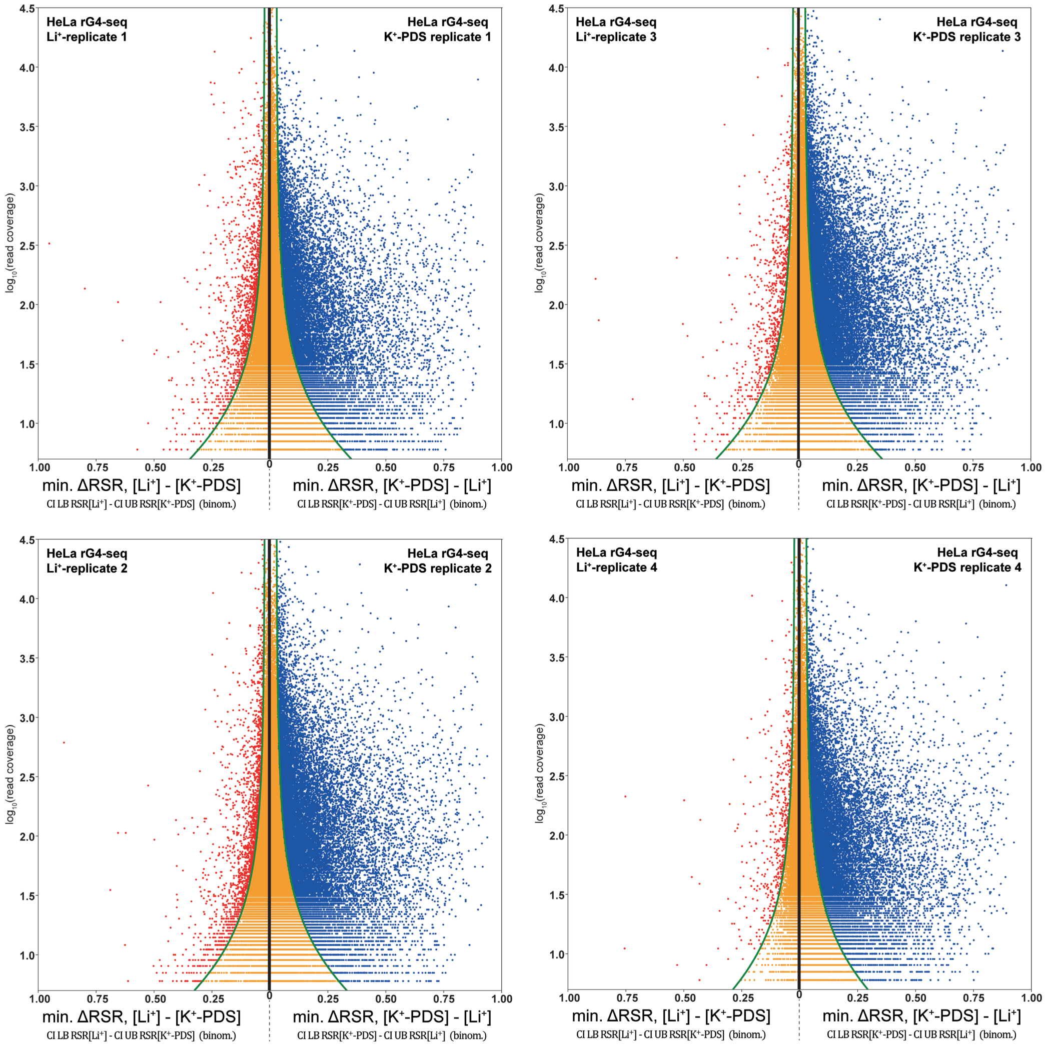

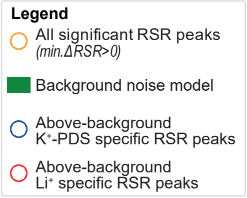

| | Alpha values for binomial proprotion confidence intervals | |
|---|---|---|
| Replicate | min. ΔRSR | Noise model |
| 1 | 0.200 | 0.130 |
| 2 | 0.200 | 0.148 |
| 3 | 0.050 | 0.120 |
| 4 | 0.200 | 0.191 |

| | No. of above-background RSR peaks | |
|---|---|---|
| Replicate | $K^+$/PDS specific | $Li^+$ specific |
| 1 | 15,035 | 1,490 |
| 2 | 22,041 | 2,087 |
| 3 | 20,717 | 833 |
| 4 | 16,768 | 545 |

### Figure S6

RSR peaks identified *ab initio* from pairwise comparison of 4 replicates of HeLa rG4-seq ($K^+$-PDS) and HeLa rG4-seq($Li^+$) by applying the minimum ΔRSR metric scheme, sequence-based filtering scheme and fragmentation-associated noise model.

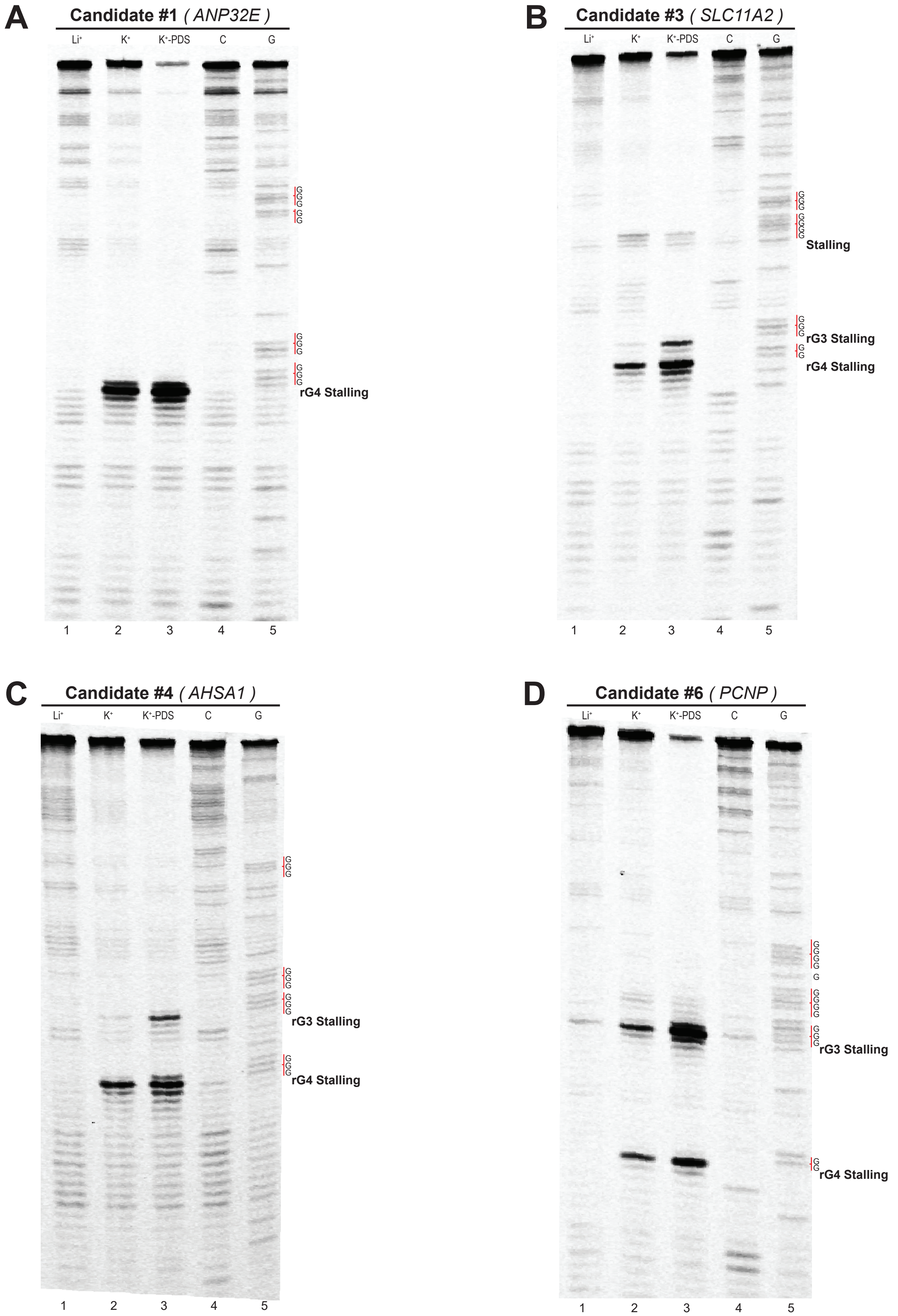

**Figure S7**

(a) RTS PAGE assay of a potential rG4 candidates on *ANP32E* gene indicated clear RTS at the 3' end of the 4th G-tract.

(b,c) RTS PAGE assay of two potential rG4 candidates on (b) *SLC11A2* gene (c) *AHSA1* gene indicated clear RTS at the 3' end of the 3rd and 4th G-tracts, respectively.

(d) A potential rG3 was initially suggested to form on *PCNP* gene based on findings from rG4-seq. RTS PAGE assay revealed a 4th G-tract 13 nt downstream of the 3rd G-tract, indicating that the region harbored a rG4 motif instead.

| Chromosome | Start | End | Strand | Gene name | 3' Sequence (30nt) | RTS site location | RTS site Contains 'GGG' Motifs | Detected by Binomial test | Detected by min. ΔRSR scheme** |
|---|---|---|---|---|---|---|---|---|---|
| **RTS site adjacent to G residues (35 sites)** | | | | | | | | | |
| chr1 | 21070287 | 21070317 | - | HP1BP3 | AGTGCTGGGATTACAGGAGTGAGCCACTGT | chr1:21070287:- | No | Y | Y |
| chr1 | 150191011 | 150191041 | - | ANP32E | GTATTTTCATGCAAATAAGTAAGGGTGGGT | chr1:150191011:- | Yes | Y | Y |
| chr1 | 180942946 | 180942976 | - | STX6 | TTACTTAAATGATTTGAGGGGTGGGAGGGA | chr1:180942946:- | Yes | Y | Y |
| chr1 | 185267080 | 185267110 | - | IVNS1ABP | GTACACTTGTGAATAAAGAGGGTGGGTGGG | chr1:185267080:- | Yes | Y | Y |
| chr1 | 229773622 | 229773652 | + | URB2 | TGCAGCAGGTTGTGCTGCAGACAGGAGCTG | chr1:229773652:+ | No | | |
| chr10 | 50732243 | 50732273 | - | ERCC6/PGBD3 | TATGAGCTGAAGCCTCTGCCCAAGGGCGGG | chr10:50732243:- | Yes | Y | |
| chr11 | 17109477 | 17109507 | - | PIK3C2A | TTGATAATATATCACTGGGACGGGGTGGGG | chr11:17109477:- | Yes | Y | Y |
| chr12 | 10758800 | 10758830 | - | MAGOHB | AGGGGTTATTTGTCATTTACAGTATTGGGG | chr12:10758800:- | Yes | Y | Y |
| chr12 | 16500625 | 16500723 | + | MGST1 | GGGAGGGCCAGGGTGGGTGGCAGATGGAAG | chr12:16500723:+ | No | | |
| chr12 | 16500635 | 16500733 | + | MGST1 | GGGTGGGTGGCAGATGGAAGACTTGGGGGG | chr12:16500733:+ | Yes | Y | |
| chr14 | 81687271 | 81687499 | - | GTF2A1 | TCTCGGCGGCGGCGGCGGCGGCGGTGGTGG | chr14:81687271:- | No | Y | |
| chr14 | 74164495 | 74164525 | + | DNAL1 | AGTTTTCTTTTCTCTCTTCTTTTCAATCTG | chr14:74164525:+ | No | | |
| chr15 | 86291231 | 86291261 | + | AKAP13 | TAGATAGACCCTGCCTTAGTAGAGGGTGGG | chr15:86291261:+ | Yes | Y | Y |
| chr16 | 19132439 | 19132469 | + | ITPRIPL2 | GGGATAAAGCATGTATAAGTTGGGAGAGGG | chr16:19132469:+ | Yes | Y | Y |
| chr16 | 85707376 | 85707406 | + | KIAA0182 | GTGACAGTCATGTGCACACATGGGCGGGGG | chr16:85707406:+ | Yes | Y | Y |
| chr17 | 3715611 | 3715641 | - | C17orf85 | TTTGGGGTTGATTTTTGTGCTGGGGTGGGA | chr17:3715611:- | Yes | Y | Y |
| chr17 | 49257909 | 49257939 | - | MBTD1 | GAGGTGGCTTAGAAACTGAGGGCGGGTGGG | chr17:49257909:- | Yes | Y | Y |
| chr17 | 60023147 | 60023177 | - | MED13 | CGATGTACAGTTTAACGGGGATAGAGGGGG | chr17:60023147:- | Yes | Y | Y |
| chr17 | 7210424 | 7210876 | + | EIF5A | CGCTTGCGCAGTGCGGGGGTGGAGGGCGGA | chr17:7210876:+ | Yes | Y | Y |
| chr17 | 40128540 | 40128570 | + | CNP | GTCCACCCCCGCCTCCCTCCCCTGCCCTCG | chr17:40128570:+ | No | | |
| chr19 | 59028606 | 59028636 | - | ZBTB45 | CTGCCCACCCCTGTGCCCCCGCCACTCGCA | chr19:59028606:- | No | | |
| chr2 | 68406110 | 68406140 | - | PPP3R1 | TTTTTTTTTTTTAGTTTGTGTGTGGGGGTG | chr2:68406110:- | Yes | | |

| | | | | | | | | | |
|---|---|---|---|---|---|---|---|---|---|
| chr2 | 201676698 | 201676923 | + | BZW1 | GGGTAGCGGCGGCCCGGGTGGGGAGGTGGG | chr2:201676923:+ | Yes | Y | Y |
| chr22 | 21356311 | 21356341 | + | THAP7-AS1 | CGGCCCGTTGTCGCCCCAACCCCGTCCCAG | chr22:21356341:+ | No | | |
| chr3 | 152017907 | 152017937 | + | MBNL1 | TGGTTGTTGCTCTTTTTTGGGGGGGTTGGG | chr3:152017937:+ | Yes | Y | Y |
| chr5 | 126172247 | 126172277 | + | LMNB1 | TTTTTTAAGTTCTTATGAGGAGGGGAGGGT | chr5:126172277:+ | Yes | Y | Y |
| chr6 | 43586704 | 43586734 | + | POLH | GGGGTTCTATATTTAGTTTGAAGAGGTGGG | chr6:43586734:+ | Yes | Y | |
| chr6 | 145172425 | 145172455 | + | UTRN | ATGTATACAGCATTGGGAAAGTGGGTGGGG | chr6:145172455:+ | Yes | Y | Y |
| chr7 | 97851053 | 97851716 | - | TECPR1 | CAGCCTCCCCTTCGCAGGTGACTGCTGGTA | chr7:97851053:- | No | | |
| chr7 | 94298466 | 94298496 | + | PEG10 | ATAATCATAAGCATTTTAGGGTGGGAGGGA | chr7:94298496:+ | Yes | Y | Y |
| chr9 | 14086231 | 14086261 | - | NFIB | CATATATATATATTCTGGGGTGGGTGGGAG | chr9:14086231:- | Yes | Y | Y |
| chr9 | 2193003 | 2193033 | + | SMARCA2 | TGGGTGGATAGTATATTTCTATGGGTGGGT | chr9:2193033:+ | Yes | Y | Y |
| chrX | 150158023 | 150158053 | + | HMGB3 | GTAACATTTTATCCAGGTTGGGGTGAGGGG | chrX:150158053:+ | Yes | Y | Y |
| **RTS site non-adjacent to G residues (7 sites)** | | | | | | | | | |
| chr1 | 202554249 | 202554279 | + | PPP1R12B | AGCAGCTAGGACTGCAGGTGTGCACCACCA | chr1:202554279:+ | No | | |
| chr15 | 56735938 | 56735968 | - | MNS1 | AAGAAACGTGAGGAGATGGAAGAAGAAAAC | chr15:56735938:- | No | | |
| chr16 | 58554140 | 58554170 | - | CNOT1 | CCTAAAGCCCACCCCTACCCTCACCCCCCC | chr16:58554140:- | No | Y | |
| chr19 | 11033127 | 11033157 | + | CARM1 | CCCCTGCAGGTCCCCCCCCGCCCCCCCACT | chr19:11033157:+ | No | | |
| chr19 | 56152393 | 56152413 | + | ZNF580 | CCTCGCACCCCCGCGCCCCC | chr19:56152413:+ | No | | |
| chr3 | 53319128 | 53319158 | - | DCP1A | TTTGTGTGCCCACCCCCACCCCCAGGGTCT | chr3:53319128:- | Yes | | |
| chr3 | 15296907 | 15296937 | + | LOC100505696 | GTCCCTCCTACCCTCATCCCCCCGCATCCC | chr3:15296937:+ | No | | |
| chr9 | 112810945 | 112810975 | + | AKAP2 | CTCCAGCGCGCCCGGAGGCTACCACTCCCT | chr9:112810975:+ | No | | |

**Table S1**

Details of 42 “Others” RTS sites previously reported in Kwok et al. (2016) and simultaneously recalled with RSR metric and binomial test / minimum ΔRSR scheme from both rG4-seq ($K^+$) and rG4-seq ($K^+$-PDS). Genomic coordinates were corresponding to GRCh37/hg19 human reference genome. Properties of the RTS sites (adjacency to G residues / containing ‘GGG’ motifs) and their recalling status were shown in the table.

| Min. Read coverage | rG4-seq condition | #Replicate | $\sum(\textit{sequenced positions})$ | $\sum(\textit{sequenced positions containing} \geq 1 \textit{ read start})$ | $P(B)$ | $\sum(\textit{read coverage of sequenced positions containing} \geq 1 \textit{ read start})$ | $\sum(\textit{read starts in sequenced positions})$ | $P(A \cap B)$ | $P(A\|B)$ |
|---|---|---|---|---|---|---|---|---|---|
| ≥6x | $K^+$ | 1 | 43,446,617 | 15,559,362 | 0.3581 | 3,565,444,556 | 63,742,568 | 0.0179 | 0.0500 |
| ≥6x | $K^+$ | 2 | 41,216,131 | 13,017,953 | 0.3158 | 3,475,638,448 | 55,074,928 | 0.0158 | 0.0500 |
| ≥6x | $K^+$ | 3 | 42,105,455 | 13,507,453 | 0.3208 | 2,938,246,465 | 53,703,833 | 0.0183 | 0.0570 |
| ≥6x | $K^+$ | 4 | 52,722,179 | 18,186,769 | 0.3450 | 4,335,282,487 | 71,534,010 | 0.0165 | 0.0478 |
| ≥6x | $K^+$-PDS | 1 | 42,869,611 | 14,025,229 | 0.3272 | 3,289,338,871 | 57,789,844 | 0.0176 | 0.0538 |
| ≥6x | $K^+$-PDS | 2 | 43,987,073 | 13,504,356 | 0.3070 | 4,094,271,905 | 60,450,721 | 0.0148 | 0.0482 |
| ≥6x | $K^+$-PDS | 3 | 45,369,082 | 14,425,694 | 0.3180 | 3,571,228,263 | 58,973,001 | 0.0165 | 0.0519 |
| ≥6x | $K^+$-PDS | 4 | 48,887,275 | 16,817,154 | 0.3440 | 3,588,040,463 | 62,777,600 | 0.0175 | 0.0509 |
| ≥6x | $Li^+$ | 1 | 51,853,328 | 18,478,171 | 0.3564 | 5,911,086,239 | 94,523,439 | 0.0160 | 0.0449 |
| ≥6x | $Li^+$ | 2 | 44,667,183 | 14,294,298 | 0.3200 | 4,498,217,619 | 68,307,246 | 0.0152 | 0.0475 |
| ≥6x | $Li^+$ | 3 | 46,364,524 | 14,839,833 | 0.3201 | 3,945,219,400 | 60,734,921 | 0.0154 | 0.0481 |
| ≥6x | $Li^+$ | 4 | 61,746,013 | 22,473,562 | 0.3640 | 6,364,155,240 | 102,464,720 | 0.0161 | 0.0442 |
| ≥2048x | $K^+$ | 1 | 205,377 | 195,888 | 0.9538 | 1,025,436,370 | 14,098,748 | 0.0137 | 0.0144 |
| ≥2048x | $K^+$ | 2 | 221,608 | 210,082 | 0.9480 | 1,135,915,016 | 13,551,271 | 0.0119 | 0.0126 |
| ≥2048x | $K^+$ | 3 | 171,093 | 161,626 | 0.9447 | 868,319,850 | 11,693,007 | 0.0135 | 0.0143 |
| ≥2048x | $K^+$ | 4 | 246,912 | 235,807 | 0.9550 | 1,263,548,976 | 16,413,174 | 0.0130 | 0.0136 |
| ≥2048x | $K^+$-PDS | 1 | 188,137 | 179,164 | 0.9523 | 960,864,841 | 12,600,469 | 0.0131 | 0.0138 |
| ≥2048x | $K^+$-PDS | 2 | 263,891 | 250,757 | 0.9502 | 1,393,577,813 | 15,607,667 | 0.0112 | 0.0118 |
| ≥2048x | $K^+$-PDS | 3 | 210,136 | 200,039 | 0.9520 | 1,103,055,047 | 13,721,284 | 0.0124 | 0.0130 |
| ≥2048x | $K^+$-PDS | 4 | 197,311 | 187,999 | 0.9528 | 978,523,049 | 13,190,069 | 0.0135 | 0.0142 |
| ≥2048x | $Li^+$ | 1 | 376,146 | 362,479 | 0.9637 | 2,057,673,637 | 26,572,792 | 0.0129 | 0.0134 |
| ≥2048x | $Li^+$ | 2 | 299,899 | 286,568 | 0.9555 | 1,615,155,435 | 19,049,791 | 0.0118 | 0.0123 |
| ≥2048x | $Li^+$ | 3 | 240,977 | 229,948 | 0.9542 | 1,308,568,932 | 15,330,434 | 0.0117 | 0.0123 |
| ≥2048x | $Li^+$ | 4 | 396,186 | 382,689 | 0.9659 | 2,082,612,795 | 27,703,361 | 0.0133 | 0.0138 |

**Table S2.**

Raw number figures for deriving the conditional probability P ( A | B ) for statistical modelling of fragmentation-associated background noise, where:

$A$: receiving 1 read start per 1x read coverage

$B$: receiving at least 1 read start

| rG4-seq condition | #Replicate | Average aligned read length | $\frac{1}{Average\ aligned\ read\ length}$ |
|---|---|---|---|
| $K^+$ | 1 | 71 | 0.014085 |
| $K^+$ | 2 | 82 | 0.012195 |
| $K^+$ | 3 | 72 | 0.013889 |
| $K^+$ | 4 | 76 | 0.013158 |
| $K^+$-PDS | 1 | 74 | 0.013514 |
| $K^+$-PDS | 2 | 87 | 0.011494 |
| $K^+$-PDS | 3 | 79 | 0.012658 |
| $K^+$-PDS | 4 | 73 | 0.013699 |
| $Li^+$ | 1 | 75 | 0.013333 |
| $Li^+$ | 2 | 82 | 0.012195 |
| $Li^+$ | 3 | 83 | 0.012048 |
| $Li^+$ | 4 | 74 | 0.013514 |

**Table S3**

Statistics of averaged aligned read lengths in HeLa rG4-seq datasets
